# Eukaryotic translation initiation factor 3d regulates stress granule assembly via its RNA binding domain

**DOI:** 10.1101/2025.11.13.688230

**Authors:** Jiaqing Lang, Aristeidis P. Sfakianos, Pablo Binder, Wei Zhang, Cathy Tournier, Mark P. Ashe, Gino B. Poulin, Alan J. Whitmarsh

**Affiliations:** Division of Molecular and Cellular Function, School of Biological Sciences, University of Manchester, Michael Smith Building, Oxford Road, Manchester, M13 9PT, UK; Division of Cancer Sciences, School of Medical Sciences, Faculty of Biology, Medicine & Health, University of Manchester, Michael Smith Building, Oxford Road, Manchester, M13 9PT, UK; Altos Labs, Cambridge, UK; Sygnature Discovery Ltd, Nottingham, UK; Turku Bioscience Centre, University of Turku and Abo Akademi University, Turku, Finland

## Abstract

Stress granules are cytoplasmic mRNA-protein complexes that form by liquid-liquid phase separation in response to a variety of stresses. Their assembly is contingent upon the inhibition of mRNA translation. Depending on the type of stress and severity, they can promote stress resistance or act to reduce cellular fitness. As such, stress granules are implicated in ageing and a range of related pathologies. Many translation factors are components of stress granules, but it is unclear how they contribute to granule assembly. Here we show that the eIF3d component of the eIF3 translation initiation complex is recruited to stress granules in human cells and is required for stress granule assembly in response to specific stresses. The RNA-binding domain of eIF3d mediates its recruitment to stress granules and deletion of this domain blocks granule formation and decreases cell viability. Furthermore, the exogenous expression of just the eIF3d RNA-binding domain can rescue stress granule assembly in eIF3d-depleted cells. We confirmed the importance of eIF3d for robust stress granule assembly *in vivo* using the nematode worm *C. elegans*. This study demonstrates that eIF3d is a critical evolutionary conserved stress granule assembly factor, rather than simply coalescing passively into stress granules following the inhibition of translation initiation.

## INTRODUCTION

Organisms need to respond to stress from their environment in order to adapt and survive. One feature of this response is a regulated change in the pattern of gene expression that can occur via both transcriptional and post-transcriptional mechanisms [1, 2]. After an acute stress, there is suppression of global mRNA translation and a shift towards the translation of specific mRNAs encoding proteins that act to mitigate the stress [2–4]. This coincides with the formation of membrane-less cytoplasmic ribonucleoprotein (RNP) granules called stress granules (SGs) that contain translationally stalled 48S pre-initiation complexes (PIC) and their associated mRNAs, together with additional RNA-binding proteins (RBPs) and other proteins [5–8]. SG assembly occurs via a process of liquid-liquid phase separation (LLPS) that is mediated by both multivalent interactions between proteins and by protein-RNA and RNA-RNA contacts [9–14]. For most stresses, the paralogous RBPs RAS GTPase-activating protein-binding protein 1 (G3BP1) and G3BP2 are essential for SG formation [15–18], but the biophysical properties and composition of SGs vary depending on the stress and cell type [7, 8, 19, 20]. SGs form rapidly and can exchange proteins and RNAs with surrounding cellular compartments throughout their lifetime, before dissolving once the stress has been resolved [9, 21–24]. SGs are generally considered to be protective and enable cells to respond and adapt to stress, with reported roles in anti-viral responses [25], inflammasome-mediated pyroptosis [26] and repairing damage to endolysosomal membranes [27]. The disruption of SG regulation can contribute to disease. For example, SGs can protect cancer cells and they can lose their transient nature and become persistent with aging, which contributes to the pathological aggregation of RBPs in neurodegenerative diseases such as amyotrophic lateral sclerosis (ALS) [19, 28, 29].

The relationship between translation inhibition, SG assembly and how this links to cell survival responses is not fully understood. Evidence suggests that SGs act as dynamic signalling hubs that regulate cell fate by coordinating stress-induced signalling with altered translation [3, 30–34]. Stresses such as oxidative stress, ER stress and viral infection require eIF2α phosphorylation-induced translation inhibition for SG assembly [19, 20], while inhibiting other translation initiation factors, for example the eIF4A RNA helicase component of the eIF4F cap-binding complex, induces SGs independently of eIF2α phosphorylation [35–37]. Many translation factors are found in SGs and several subunits of the eIF3 translation initiation complex have been proposed as core components of the SG interaction network that contributes to SG assembly [5, 7, 8, 18, 38], but the mechanisms involved are poorly characterised.

The eIF3 complex is around 800 kDa and is made up of twelve subunits in humans and assembles into two subcomplexes: the octamer containing eIF3a, c, e, f, h, k, l, and m, and the yeast-like core (YLC) containing eIF3b, g and i [39]. eIF3d is not a core component of the eIF3 complex but is recruited to the octamer via eIF3e [40, 41]. During canonical 7-methyl guanosine (m^7^G) cap-dependent mRNA translation initiation, the eIF3 complex coordinates multiple steps in forming the 43S PIC and critically links it to the 5’ end of mRNA via interactions between the eIF4F complex bound at the mRNA cap and the eIF3c, d and e subunits on the 43S PIC [42–44]. In addition, eIF3 takes part in 43S PIC scanning of the 5’ UTR for the AUG start codon and likely persists on the ribosome during early elongation [45, 46]. eIF3 also regulates translation through alternative mechanisms. It may be recruited directly to the 5’ cap independently of the eIF4F complex via the eIF3d subunit and, in collaboration with eIF4G2/DAP5, mediates translation of subsets of mRNAs that contribute to stress survival, T-cell regulation and cell migration [47–52].

To start to address how eIF3 factors contribute to SG biology, we focused on eIF3d. It is an important mediator of the translational response to stress [48, 50, 53] and two genetic screens in the human osteosarcoma cell line U2OS have implicated it in SG formation in response to sodium arsenite and heat stress [18, 38]. Furthermore, the knockdown of eIF3d does not affect the formation of the core eIF3 complex [39, 54]. We therefore set out to uncover the molecular determinants of eIF3d recruitment to SGs and whether it has roles in SG assembly independently of the core eIF3 complex.

## RESULTS

### eIF3d is required for SG assembly in HeLa and U2OS cells in response to specific stress conditions

To obtain a broad picture of the role of eIF3d in SG formation in response to different stresses, we knocked-down the expression of eIF3d in human cells using siRNA. Two siRNAs were used to target eIF3d in HeLa cells and successful knock-down was confirmed by immunoblot (Suppl Fig. 1A). The cells were subjected to sodium arsenite (50 μM, 1 h), heat stress (43°C, 1 h), ER stress (20 μM thapsigargin, 1 h) or osmotic stress (0.2 M NaCl, 1 h). eIF3d colocalised with the SG marker protein G3BP1 under all conditions indicating its recruitment to SGs (Fig. 1). The knock-down of eIF3d by both siRNAs strongly suppressed SG formation in response to sodium arsenite stress, heat stress and ER stress based on the loss of G3BP1 positive granules, but not in response to osmotic stress (Fig. 1; Suppl Fig. 2). A similar pattern of responses was observed in a second cell line, U2OS (Suppl Figs. 1B and 3). Of note, a higher dose (100 μM) of sodium arsenite was required to robustly induce SGs in U2OS compared to HeLa indicating differential sensitivities to arsenite stress between the cell lines. Arsenite stress, heat stress and ER stress induce translational repression and SG assembly via the phosphorylation of eIF2α, while osmotic stress is reported to promote SGs independently of this [16, 19, 20, 55]. Another mechanism of translation inhibition which leads to SG formation independently of phosphorylated eIF2α is the targeted inhibition of the RNA helicase eIF4A [20, 35, 36]. We therefore treated HeLa cells with the eIF4A inhibitor FL3 (0.5 μM, 24 h) [56, 57]. eIF3d was present in FL3-induced SGs, but its knock-down did not affect their formation (Suppl Fig. 4). In summary, eIF3d is required for SG assembly after moderate arsenite stress, heat stress and ER stress, but not in response to osmotic stress or eIF4A inhibition.

**Figure 1.**
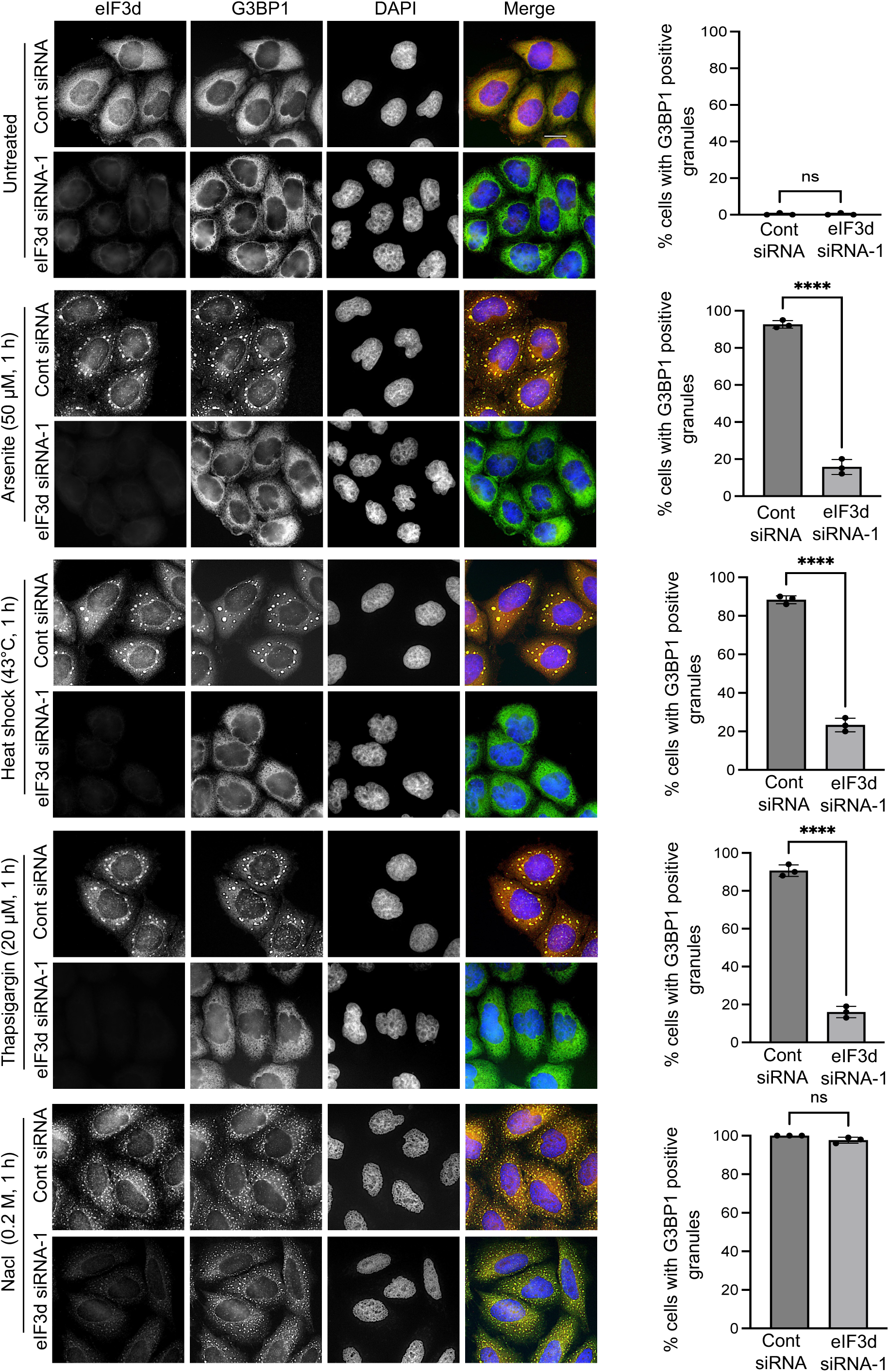
eIF3d is selectively required for SG formation induced by different stress. HeLa cells were transfected with either control (Cont) siRNA or eIF3d siRNA-1 for 48 h before treatment with the indicated stresses. Cells were probed with antibodies against eIF3d and G3BP1. Nuclei were stained with DAPI. Representative images of cells are shown in the left-hand panels. Scale bar = 20 μm. Quantification of cells with G3BP1-positive granules is shown in the right-hand panels. A total of 100 cells were analysed in each of three biological replicates. Error bars represent standard deviation. Statistical analysis was performed using unpaired t-tests (*ns* = not significant, ****P < 0.0001).

### The RNA binding domain of eIF3d is required for its targeting to SGs

We next sought to determine how eIF3d is recruited to SGs. The eIF3d domain structure features an N-terminal region with binding sites for eIF3e and other eIF3 subunits [41, 58, 59], an RNA binding domain (RBD) [60], the proposed cap-binding region [49], plus a C-terminal acidic region of unknown function (Fig. 2A). We initially generated two deletion mutants, one containing the N-terminus (amino acids 1-114) that includes the eIF3 binding region and the RBD, and one covering the rest of the protein, including the cap-binding domain and the acidic tail (amino acids 115-548) (Fig. 2A). These protein fragments can occur due to cleavage of eIF3d by the HIV protease [61]. When expressed in HeLa cells subjected to arsenite-induced stress, the eIF3d 1-114 fragment localised to SGs but the 115-548 fragment did not (Fig. 2B). This indicated that the key determinants of eIF3d localisation to SGs reside in the N-terminal part of the protein containing the contact sites with the core eIF3 complex and the RBD. As protein-RNA contacts are an essential feature of RNP condensates [9, 12, 13], we generated an eIF3d deletion mutant lacking the RBD, eIF3d(ΔRBD) (Fig. 2A, C). This mutant did not localise to SGs following sodium arsenite stress, heat stress or thapsigargin-induced ER stress in HeLa cells (Figs. 2D, E; Suppl Fig. 5). Previous studies have implicated Arg residues within RBPs in promoting phase separation [62]. As the eIF3d RBD contains several Arg residues, we mutated five of these to Lys residues and found that this mutant, eIF3d(5RK), was also excluded from SGs in response to sodium arsenite stress, heat stress and ER stress (Fig. 2; Suppl Fig. 5). Similarly, the loss or mutation of the RBD blocked eIF3d recruitment to SGs in U2OS cells (Suppl Fig. 6). We conclude that the RBD is the major determinant of eIF3d recruitment to SGs.

**Figure 2.**
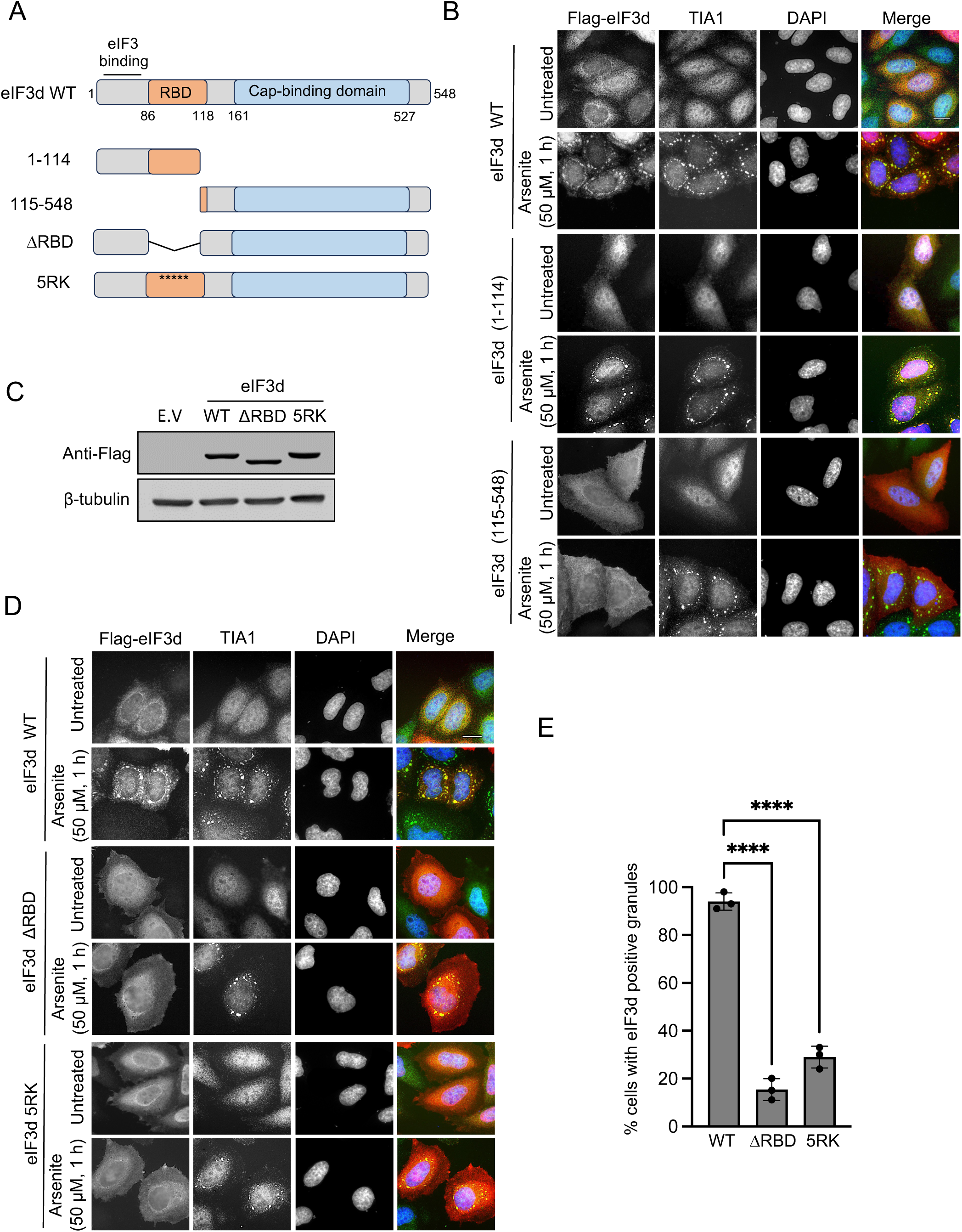
The RNA binding domain (RBD) of eIF3d is required for its recruitment to SGs. **A.** Schematic representation of the domain structure of eIF3d and the mutants generated. RBD = RNA-binding domain. 5RK = five arginine residues mutated to lysine within the RBD. **B.** HeLa cells were transfected with plasmids expressing Flag-tagged eIF3d (WT) or the truncated eIF3d mutants: 1–114 (containing the RBD) and 115–548 (lacking the RBD). Cells were treated with 50 µM sodium arsenite for 1 h. TIA1 immunofluorescence was used to visualise stress granules (SGs). Scale bar = 20 µm. **C.** Protein expression levels of the WT and the ΔRBD and 5RK mutants were assessed by immunoblotting with anti-Flag antibody. β-tubulin was used as a control to ensure equal protein input. **D-E.** HeLa cells were transfected with plasmids expressing Flag-tagged eIF3d wild-type (WT), ΔRBD, or 5RK mutants and treated with 50 µM sodium arsenite for 1 h. TIA1 immunofluorescence was used for SG visualisation. Scale bar = 20 µm. Representative images of cells are shown in D and quantification of the percentage of cells with eIF3d-positive granules is shown in E. A total of 100 cells were analysed in each of three biological replicates. Error bars represent standard deviation. Data were analysed using one-way ANOVA (****P < 0.0001).

### The RNA binding domain of eIF3d is required and sufficient for SG formation

While the RBD is critical for targeting eIF3d to SGs, it was important to determine if it was also required for SG assembly. To examine this, we depleted endogenous eIF3d from HeLa cells with siRNA and exogenously expressed wild-type eIF3d or the ΔRBD mutant. The expression of wild-type eIF3d, but not the ΔRBD mutant, rescued the suppression of arsenite-induced SG assembly caused by endogenous eIF3d knock-down. This demonstrated the requirement of the RBD for eIF3d to promote SG assembly (Fig. 3). We next asked the question whether the RBD alone was sufficient to be targeted to SGs and facilitate their assembly. We fused the RBD with the red fluorescent protein mCherry and found that it was predominantly nuclear. However, we did observe weak recruitment of the RBD-mCherry fusion protein to cytoplasmic SGs (Suppl Fig. 7). To avoid the complication of having a limited pool of cytoplasmic RBD-mCherry protein available for incorporation into SGs, we tagged the RBD with the strong nuclear export signal (NES) from PKI [63] (Fig. 4A). This fusion protein displayed increased cytoplasmic localisation and was robustly recruited to SGs in response to arsenite and heat stress (Fig. 4B). The expression of the NES-tagged RBD fused to mCherry significantly rescued SG formation in cells depleted of endogenous eIF3d, with over 50% of cells displaying SGs compared to 80% of cells that expressed the full-length eIF3d (Fig. 4C and D). The NES-tagged RBD mutant with five Arg mutated to Lys (5RK) could not rescue SG formation in the eIF3d-depleted cells (Fig. 4C and D). We conclude that the RBD of eIF3d is required for it to coalesce into SGs and that it can act as an autonomous SG targeting domain that it is sufficient to support SG assembly.

**Figure 3.**
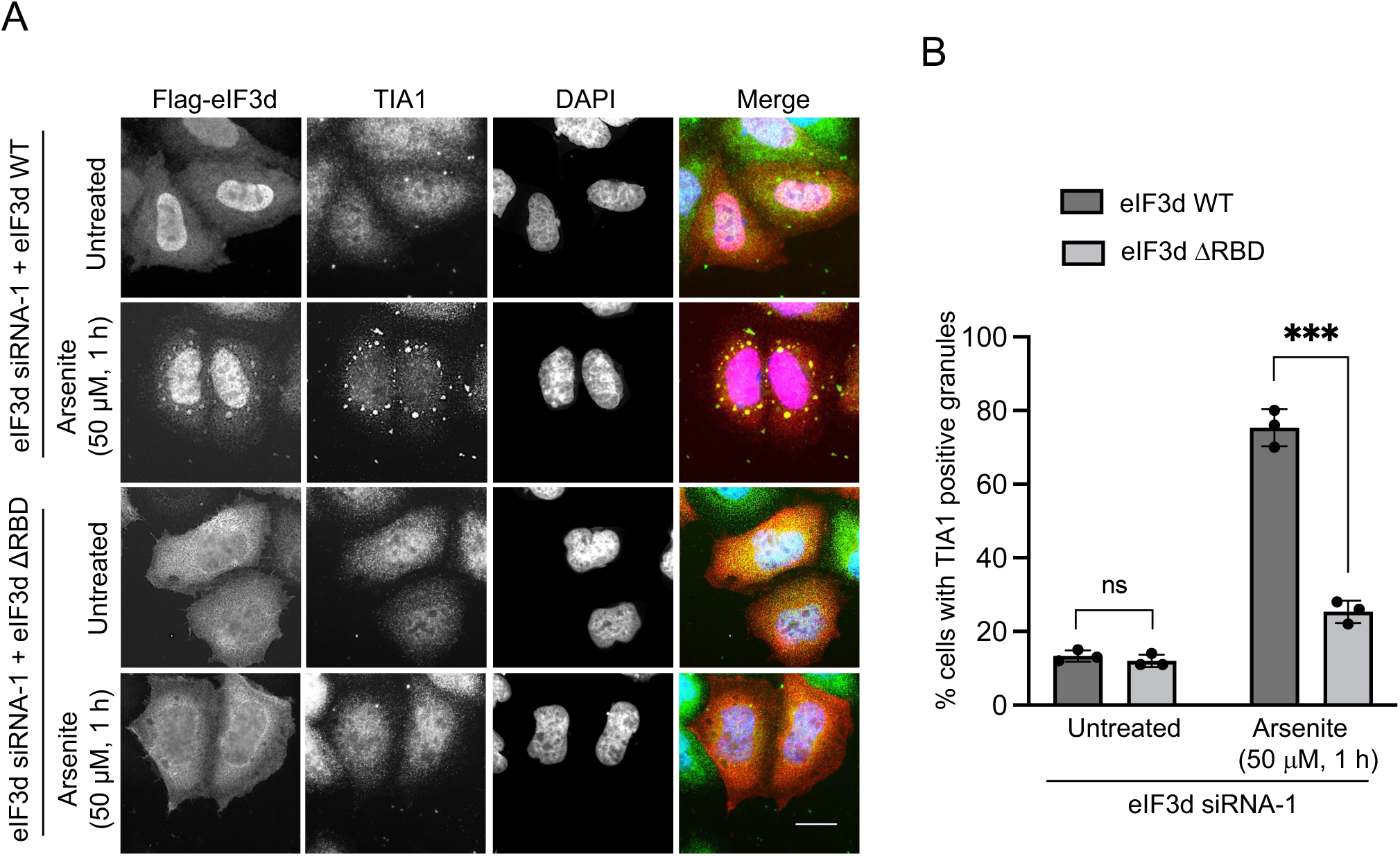
The RBD is required for eIF3d to support SG assembly. **A.** HeLa cells were transfected with eIF3d siRNA and either a plasmid expressing Flag-tagged wild-type eIF3d (WT) or eIF3d ΔRBD. Cells were then left untreated or treated with sodium arsenite (50 μM, 1h) and probed with antibodies against the Flag-tag and TIA1. Scale bar = 20 µm. **B**. Quantification of the percentage of cells with TIA1-positive granules. A total of 100 cells were analysed in each of three biological replicates. Error bars represent standard deviation. Data were analysed using unpaired t-tests (*ns* = not significant, ***P <0.0005).

**Figure 4.**
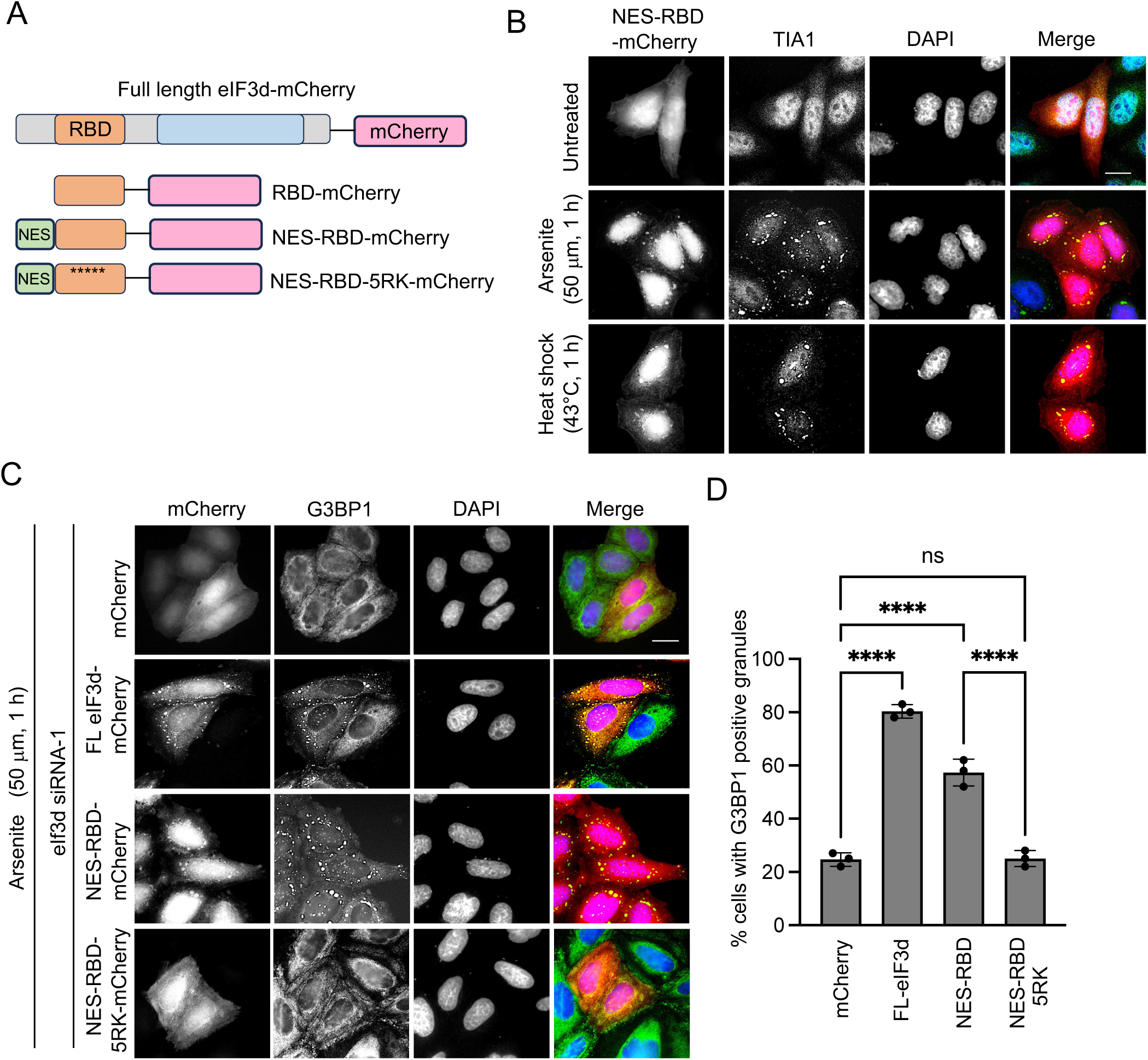
The RBD of eIF3d can act as an autonomous SG targeting domain and support SG assembly. **A.** Schematic representation of the eIF3d-mCherry fusion mutants generated. RBD = RNA-binding domain. 5RK = five arginine residues mutated to lysine within the RBD. NES = the nuclear export sequence from PKI. **B**. HeLa cells were transfected with a plasmid expressing the NES-RBD fused to mCherry. Cells were then treated with either 50 µM sodium arsenite or heat stress at 43 °C for 1 h. TIA1 immunofluorescence was used to visualise SGs. Scale bar = 20 µm. **C**. HeLa cells were transfected with eIF3d siRNA and plasmids expressing either mCherry alone, full-length (FL) eIF3d fused to mCherry, the NES-RBD fused to mCherry, or the NES-RBD-5RK mutant fused to mCherry. Cells were then treated with 50 µM sodium arsenite for 1 h. Scale bar = 20 µm. **D**. Quantification of cells with G3BP1-positive granules for the indicated mutants. A total of 100 cells were analysed in each of three biological replicates. Error bars represent the standard deviation. Data was analysed using one-way ANOVA (*ns* = not significant, ****P < 0.0001).

### The deletion of the eIF3d RBD impairs translation and cell survival

To further test the functional importance of the eIF3d RBD, we established stable HeLa cell lines featuring doxycycline-inducible shRNA knock-down of endogenous eIF3d and re-expression of wild-type or mutant eIF3d. In agreement with our previous siRNA data (Fig. 3), the induction of shRNA expression greatly reduced arsenite-induced SG assembly, which was rescued by wild-type eIF3d but not the ΔRBD mutant (Fig. 5A and B; Suppl Figs. 8A and B). We also investigated the second RNA interacting domain of eIF3d, the cap-binding domain. Mutation of key residues required for cap-binding [49] did not affect eIF3d localisation to SGs or SG assembly (Suppl Fig. 8C). These results confirmed that the RBD of eIF3d is the key domain required to promote SG assembly. To address the functional consequences of this, we performed a puromycin-incorporation assay to determine the importance of the eIF3d RBD for global translation. The knock-down of eIF3d reduced global translation by around 50% and this was rescued by wild-type eIF3d but not by the mutant lacking the RBD (Fig. 5C). The mutation of the cap-binding region did not significantly impact global translation levels, supporting the evidence that the cap-binding function of eIF3d is directed towards highly specific subsets of mRNAs [47, 49, 50–52]. We next determined whether the eIF3d RBD was important for cell survival in response to arsenite stress. The knock-down of eIF3d reduced cell number by about 70% in response to prolonged sodium arsenite stress (30 μM, 14 h) indicating the importance of eIF3d for cell viability (Fig. 5D). The expression of either wild-type or the cap-mutant eIF3d rescued this decrease in cell number, but the ΔRBD mutant did not (Fig. 5D). Taken together, our data show that the RBD domain of eIF3d is critical for its role in SG formation and cell viability in response to arsenite stress.

**Figure 5.**
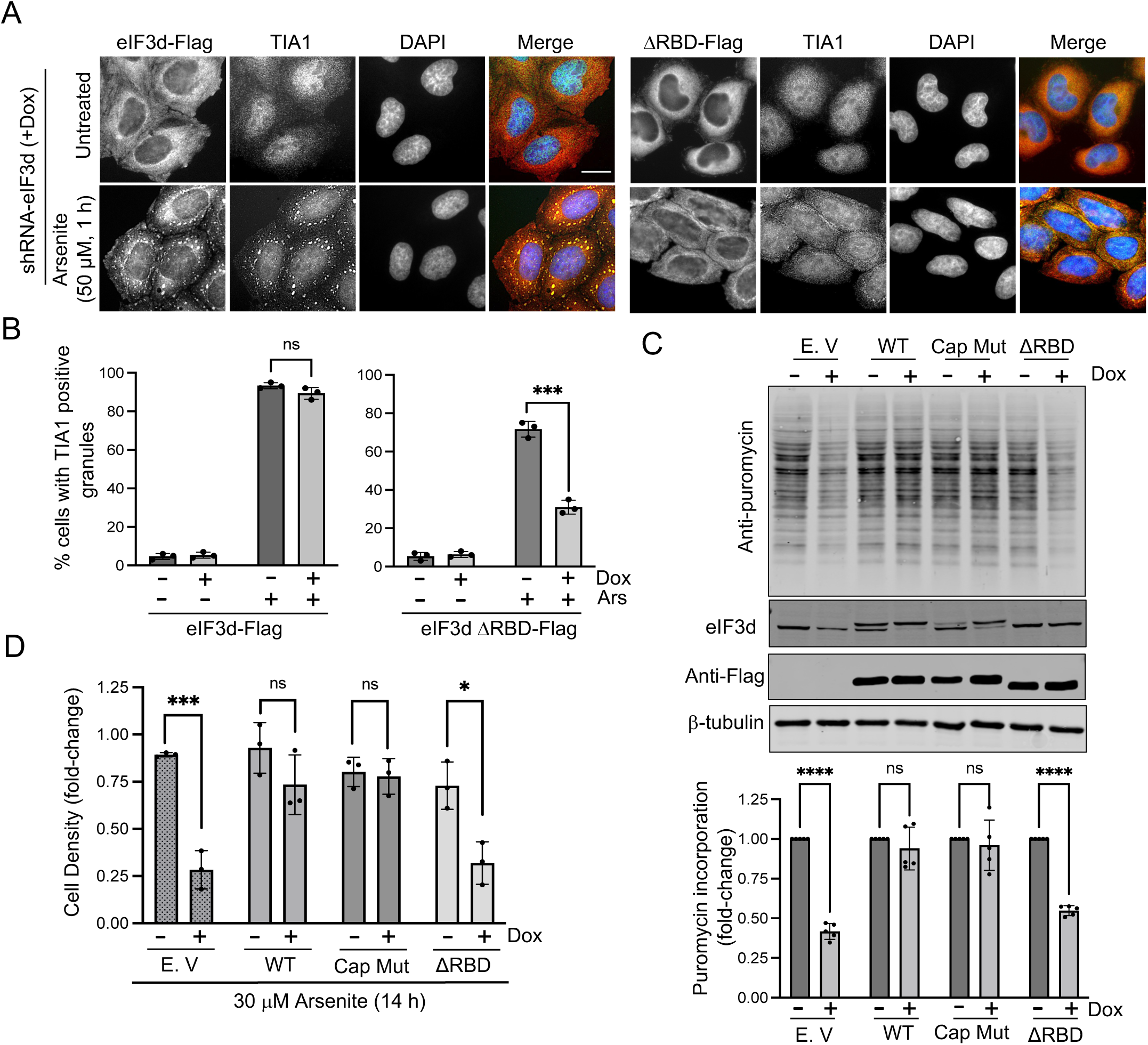
The eIF3d RBD is required for *de novo* protein synthesis and cell viability in response to arsenite stress. Stable cell lines expressing an inducible shRNA targeting endogenous eIF3d and either empty vector (E.V), Flag-eIF3d WT, Flag-eIF3d ΔRBD or Flag-eIF3d Cap Mut were established. **A**, **B**. The eIF3d shRNA was induced with 100 ng/ml doxycycline for 48 h before treatment with 50 μM sodium arsenite for 1 h. TIA1 immunofluorescence was used for SG visualisation. Flag-eIF3d WT and Flag-eIF3d ΔRBD were detected with an anti-Flag-tag antibody. Scale bar = 20µm. Representative images of cells are shown in A and quantification of cells with TIA1-positive granules under the indicated conditions is shown in B. 100 cells were analysed in each of 3 biological repeats. Error bars represent the standard deviation. Data was analysed using two-sample t-test (ns = not significant, ***P<0.0002). **C.** The Flag-eIF3d, Flag-eIF3d ΔRBD and Flag-eIF3d Cap Mut stable cell lines were induced with 100 ng/ml doxycycline for 48 h to knock down endogenous eIF3d expression. Incorporation of puromycin into newly synthesised proteins was assessed by immunoblotting. Lysates were also probed with antibodies against eIF3d and the Flag epitope. Band intensities from five independent experiments were quantified and normalised to β-tubulin levels. Quantification of newly synthesised protein is presented as fold change relative to the -Dox controls. Data were analysed using t-tests (ns = not significant; ****P <0.0001). **D.** The stable cell lines were induced with 100 ng/ml doxycycline for 48 h to knock down endogenous eIF3d expression and then treated with 30 µM sodium arsenite for 14 h. Cell density was quantified. Error bars represent the standard deviation. Data were analysed using t-tests (ns = not significant; *P <0.015, ***P <0.0005).

### The eIF3d orthologue EIF-3.D is required for stress granule formation in *C. elegans*

Having established the importance of eIF3d for SG assembly in cultured human cells, we addressed if this was also the case in an animal model. The stress pathways regulating translation and the components of the translation initiation machinery are evolutionary conserved between humans and the nematode worm *C. elegans* [64]. We therefore employed two SG reporter strains combined with RNAi knockdown of the worm eIF3d orthologue EIF.3.D. The reporters feature pharyngeal expression of either red fluorescent protein (RFP) fused to the stress granule associated RNA binding protein PAB-1 (orthologue of human PABPC1) or Venus fluorescent protein fused to TIAR-2 (an orthologue of human TIA1/TIAR) [65]. Subjecting the reporter worms to heat stress (36°C, 3 h) induced the formation of RFP-PAB-1 and Venus-TIAR-2 positive granules in the pharynx (Suppl Fig. 9A). When the reporter worms were fed with *E. coli* expressing *eif-3.D* RNAi, SG formation was severely diminished (Fig. 6). We conclude that EIF-3.D is required for heat-induced SG formation in *C. elegans*. To gain a broader picture of the importance of eIF3 for SG formation *in vivo*, we extended our RNAi analysis to the other subunits of the complex. We found that RNAi knock-down of multiple eIF3 subunits reduced granule formation to varying degrees (Suppl Figs. 9B and C). It was noted that the knock-down of some of these factors caused severe developmental defects (i.e. for EIF-3.B and G). In contrast, the knock-down of EIF-3.D and other factors, including EIF3.E and EIF-3.F, strongly impaired granule formation without having a drastic effect on development (Suppl Figs. 9 and 10). These data suggest that, in addition to eIF3d, other eIF3 factors also play important roles in SG assembly *in vivo*.

**Figure 6.**
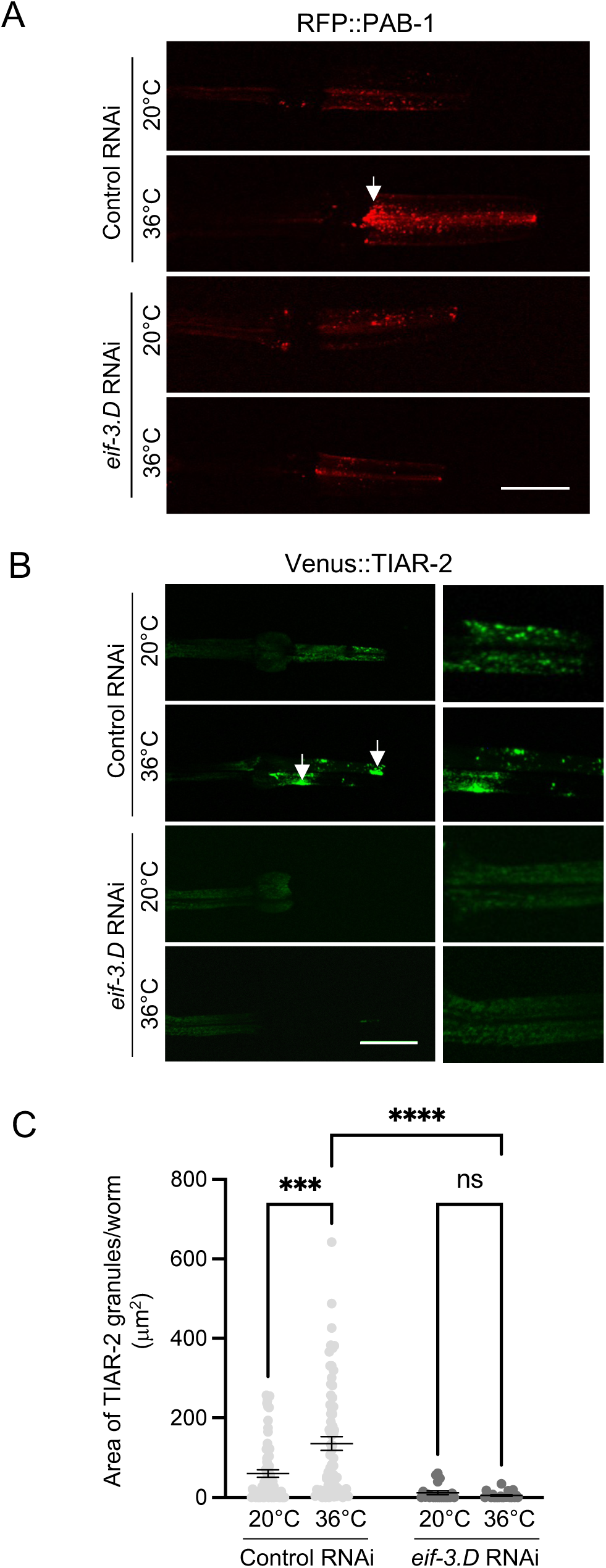
*C. elegans* EIF-3.D is required for stress granule formation in vivo. **A, B.** Worms expressing pharyngeal RFP::PAB-1 and Venus::TIAR-2 reporters were fed *eif-3.D* RNAi and subjected to heat shock at 36 °C for 3 h. Representative confocal images of the pharynx are shown for the indicated conditions. The right-hand panel in B shows a magnified area. Scale bar = 25 μm. **C.** Quantification of the total area of TIAR-2–positive granules per worm for each of the indicated conditions. At least 30 worms were analysed per condition across three biological replicates. Error bars represent s.e.m. Data were analysed by two-way ANOVA (*ns* = not significant; ***p<0.0005; ****p < 0.0001).

## DISCUSSION

The eIF3 complex is essential for the initiation of canonical cap-dependent translation in collaboration with eIF4F and also mediates alternative mechanisms of translation initiation [42, 43, 53]. In this study, we find that eIF3d is required for SG assembly in both human cell lines and in *C. elegans* (Figs. 1 and 6) and that the RBD of eIF3d is essential for this (Figs. 4 and 5). This establishes eIF3d as an important SG regulatory factor. Our findings support experiments by Yang et al [18] who built on previous proteomic and functional screens [5, 38] to define a core network of 36 proteins that are localised to SGs and are important for their assembly. These proteins include eIF3d, alongside well-characterised SG regulators such as G3BP1, G3BP2, TIAR, TIA1 and UBAP2L [18]. Additional eIF3 factors are also part of this network including eIF3g, eIF3i and eIF3e [18]. Our *C. elegans* RNAi screen also identified these factors as important SG regulators *in vivo* (Suppl Figs. 9 and 10). This suggests that eIF3d along with other eIF3 factors and a subset of RBPs play a central role in the coalescence of translationally stalled 48S pre-initiation complexes and their associated mRNAs into SGs in response to specific stresses.

Structural analysis of translation initiation complexes indicates that eIF3d forms part of the mRNA exit channel making contacts with eIF3e, eIF3a and eIF3c, as well as associating with the eIF4G component of the eIF4F complex and contacting 40S subunits and 18S rRNA [40, 44, 59, 66], The RBD is clearly important for eIF3d function in translation as the ΔRBD mutant fails to rescue the decreased translation observed in eIF3d-depleted cells (Fig. 5C). An important question is whether eIF3d has a distinct role in SG assembly or is acting within the context of the eIF3 complex or translational initiation complexes. The RBD does not include either the region of eIF3d that binds to eIF3e or the eIF3c and eIF4G contact sites [40, 44, 59, 66]. Therefore, the finding that it acts as an autonomous SG recruiting domain and supports SG assembly when fused to mCherry (Fig. 4) argues that the major function of eIF3d in SG assembly is independent from the rest of the eIF3 complex. However, the rescue of SG assembly in eIF3d knock-down cells is less robust when expressing the RBD alone compared to full-length eIF3d (Fig. 4). This suggests that, while the RBD is essential for supporting SG assembly, eIF3d interactions with the eIF3 complex or other translation factors, potentially within the context of the stalled 48S PIC, may have contributory roles.

How the RBD promotes eIF3d recruitment to SGs and their formation is unclear. The mutation of Arg residues to Lys within the RBD prevents its localisation to SGs and fails to support SG assembly (Figs. 2D and E; Figs. 4C and D). Arg residues within RBDs can mediate multiple interactions with RNA through both hydrogen bonding and electrostatic interactions. It is possible that eIF3d is recruited to SGs via interactions with RNA and helps to nucleate and stabilise the condensates. Alternatively, or in combination, the RBD may form contacts with other proteins. Previous studies have identified multivalent interactions between Arg and aromatic side chains such as Tyr as drivers of phase separation [62]. Interestingly, the Arg residues in the eIF3d RBD can be methylated [67, 68]. Arginine methylation within RBPs is reported to usually reduce their propensity to phase separate [69], although may promote phase separation in specific contexts [70]. It will be interesting to determine if Arg methylation of the eIF3d RBD is a key modification regulating its functions during stress.

An important aspect of our study is that the requirement for eIF3d in SG formation is stress-specific. Stresses that induce canonical SGs downstream of eIF2α Ser-51 phosphorylation, including heat, moderate sodium arsenite stress and ER stress, require eIF3d for SG assembly (Fig. 1; Suppl Figs 2 and 3). However, eIF3d is not required for the assembly of non-canonical SGs that form independently of eIF2α phosphorylation following eIF4A inhibition or osmotic stress, although it still localises to these granules (Fig. 1; Suppl Figs. 2 - 4). This selective requirement for eIF3d may reflect differences in the importance of specific proteins or the prominence of RNA-RNA interactions depending on the type of stress and how it impacts translation. For example, it has been shown that SGs formed after eIF4A inhibition are more reliant on RNA-RNA interactions, reflecting the proposed role of eIF4A as an RNA chaperone that limits intermolecular RNA interactions [37]. It is less clear how osmotic-stress induced SGs form as they are independent of both phosphorylated eIF2α and G3BP1 [16, 20, 55]. One proposed mechanism is through a combination of macromolecular crowding due to cell shrinkage, combined with the increased ionic strength reducing electrostatic repulsion between mRNAs [71]. This suggests that eIF3d may only become essential for SG assembly under conditions where RNA-RNA interactions are suppressed by eIF4A or physical and chemical changes to cells increase the likelihood for proteins and RNAs to coalesce.

A remaining question is how eIF3d functions are integrated in response to stress. As we have shown, a pool of eIF3d is recruited to SGs and is required for their assembly. However, when canonical eIF4E-dependent translation initiation is suppressed, eIF3d binds to the 5’ cap of stress-specific mRNAs and mediates their translation [47–52]. While translation has been reported to occur in the vicinity of SGs [72], there is little evidence of this being widespread, suggesting that most stress-specific translation is occurring on ribosomes in the cytoplasm. Therefore, there may be distinct pools of eIF3d under stress conditions, one involved in the translation of essential stress-specific transcripts and one that participates in SG assembly. Furthermore, we also detect endogenous eIF3d in the nucleus (Fig. 1; Suppl Figs. 2 and 3) and the eIF3d RBD by itself is strongly nuclear suggesting that it may have a broader role in regulating eIF3d localisation (Suppl Fig. 7). The function of nuclear eIF3d is unclear, although a role in RNA splicing has been proposed [73]. The latter may also link to eIF3d’s role in SGs as other splicing regulators are also core SG components [5, 7, 18]. It will be important to determine how these different pools of eIF3d communicate and integrate as part of the cellular stress response.

It is reported that eIF3d expression is upregulated in many cancers and may promote cell proliferation and survival of some cancer types [53]. While there is evidence to suggest that this may be via regulating the translation of cell-cycle regulators, it is possible that high levels of eIF3d promote SG formation in cancer cells, thus contributing to their survival and resistance to anti-cancer drugs. Previous studies have implicated other core SG proteins such as G3BP1, CAPRIN-1 and TIA1 in promoting hallmarks of cancer [29]. Therefore, manipulating core SG proteins, such as eIF3d, may limit tumourigenesis and enhance anti-cancer therapies.

In conclusion, we have characterised the function of the translation initiation factor eIF3d in SG assembly, adding to its expanding functional repertoire and supporting the evidence that it is a key mediator protecting cells from stress. Our data also suggest a more extensive role for eIF3 complex components in SG regulation and it will be interesting in future to determine their precise mechanisms, whether as part of eIF3 subcomplexes or acting independently.

## MATERIALS AND METHODS

### Cell culture and treatments

U2OS and HEK293T were purchased from American Type Culture Collection (ATCC) and cultured in Dulbecco’s Modified Eagles Medium (D6429, Sigma-Aldrich). HeLa M cells were cultured in Dulbecco’s Modified Eagles Medium (D5796, Sigma-Aldrich). Both media were supplemented with 10% feotal bovine serum (FBS) (10500064, ThermoFisher). Cells were grown at 37°C in 5% CO_2_. Stress granules were induced by addition of NaASO_2_ (S7400, Sigma-Aldrich) or Thapsigargin (CAY10522, Cambridge Bioscience) at the concentrations and times indicated in the figure legends. Heat stress was applied by incubating cells with 43°C in 5% CO_2_ for 1 hr. Mycoplasma testing was routinely performed every month as described [74].

### Transfection of plasmids and siRNAs

Plasmids were transfected into cells using JetPEI (101B-010N, Polyplus Transfections). Cells were incubated 6 hr in the presence of plasmid/JetPEI followed by changing the medium and incubation overnight prior to analysis. Plasmid used were pCDNA3-Flag-eIF3d, pCDNA3-Flag-eIF3d (1-114), pCDNA3-Flag-eIF3d (115-548), pCDNA3-Flag-eIF3d (ΔRBD) featuring a deletion of amino acids 86 to 118, pCDNA3-Flag-eIF3d (5RK) featuring arginine residues 95, 97, 99, 103 and 106 all changed to lysine, pmCherry-N1-eIF3d, pmCherry-N1-RBD, pmCherry-N1-PKI-NES-RBD and pmCherry-N1-PKI-NES-RBD (5RK). Details of plasmid generation will be provided upon request. siRNAs were transfected into cells plated in 6 well dishes at a concentration of 20nM using Lipofectamine RNAiMAX transfection reagent (13778150, ThermoFisher Scientific). Control siRNA (5’-AGGUAGUGUAAUCGCCUUG-3’) was made by Eurofins Technologies and siRNAs targeting the *EIF3D* 3’UTR (Hs_EIF3D_1 SI05097344, Hs_EIF3D_7 SI04273612) were purchased from Qiagen.

### Generation of stable cell lines

To generate doxycycline inducible shRNAs, double-stranded oligonucleotides targeting the 3’ UTR of *EIF3D* transcripts (Fwd: 5’-CCGGCTTAGTGGAATGTGTGTCTAACTCGAGTTAGACACACATTCCACTAAGTTTTTG-3’; Rev: 5’-AATTCAAAAACTTAGTGGAATGTGTGTCTAACTCGAGTTA GAC ACA CATTCC ACTAAG-3’) and non-targeting control (Fwd: 5’-CCGGCCTAAGGTTAAGTCGCCCTCGCTCGAGCGAGGGCGACTTAACCTTAGGTTTTTG-3’; Rev: 5’-AATTCAAAAACCTAAGGTTAAGTCGCCCTCGCTCGAGCGAGGGCGACTTAACCTTAGG-3’) were synthesised by Eurofins Technologies and subcloned into Age I and Eco RI restriction enzymes in Tet-pLKO-puro lentiviral plasmid (Addgene #21915). The lentiviral plasmid expressing wild-type eIF3d fused to 3xFlag sequence under the control of the hPGK promoter was designed by the Whitmarsh Lab and purchased from VectorBuilder. The eIF3d(ΔRBD) mutant, eIF3d Cap Mutant (D249Q/V262I/Y263A) and empty vector control were constructed using Q5 site-directed mutagenesis kit (E0554; NEB). Lentiviral particles were generated with HEK293T cells as previously described [75]. Infected cells were selected by incubation with 2 µg/mL puromycin and 10 µg /ml blasticidin until no live cells remained in the non-infected group. Resistant colonies were pooled and expanded in antibiotic-free medium.

### Protein extraction and immunoblot analysis

Cells were washed with PBS and lysed in RIPA buffer (R-0278, Sigma-Aldrich) supplemented with protease and phosphatase inhibitors (87786 and 78420, ThermoFisher Scientific). Protein concentrations were quantified by DC^TM^ protein assay (500-0112, Bio-Rad). Protein lysates were loaded on 10% or 14% polyacrylamide gels (A2-0072, ProtoGel) and electrophoresis performed at 100 V for 2 hr. Resolved proteins were transferred to nitrocellulose (NC) membrane (1704270, Biorad transfer kit) using Trans-Blot Turbo Transfer System (25V 2.5mA 7min). Membranes were blocked with 1x casein blocking buffer (86429, Sigma-Aldrich) in TBS (J60877.K2, Thermo Scientific) for 1 hr at room temperature and incubated with primary antibodies in TBST (28360, Thermo Scientific) overnight at 4 °C. Membranes were washed in TBST buffer and incubated for 1 hr at room temperature with IRDye® 800CW anti-rabbit or anti-mouse secondary antibodies (LI-COR) diluted in casein/TBST, prior to washing with TBST buffer and imaging using the Odyssey infrared system (LI-COR Biosciences). Fluorescent signals were quantified using Empiria studio v1.1. Antibodies used are described in Supplementary Table 1. Unprocessed scans of immunoblots are provided in Supplementary Figure 11.

### Immunofluorescence microscopy

Cells were grown on cover slips and fixed in 4% Paraformaldehyde (158127, Sigma-Aldrich) for 15min at room temperature. Cells were permeabilised with PBS (60-00010-11, PluriSelect) containing 0.2% Triton X-100 for 20min and blocked with 3% BSA (5217, Tocris) in 0.1% Triton X-100 PBST for 30min. Cover slips were incubated at 4°C with primary antibodies to eIF3d, G3BP1, TIA1 or Anti-Flag M2 (see Suppl Table 1). Following washes with 0.1% PBST, cells were incubated in 3% BSA in 0.1% PBST solution containing Alexa Fluor™ fluorescent secondary antibodies (Invitrogen; see Suppl Table 1) for 1h at room temperature, before further washes in PBST and mounting using the Prolong® Diamond-DAPI (P36962, ThermoFisher). Cover slips were sealed onto slides with antifade coverslip sealant (23005, Biotium) after mounting overnight. Imaging was performed on a Leica DM500B Fluorescence microscope with filters set for DAPI, FITC and Texas Red. Images were collected with a Leica DCF340 FX camera at a magnification of 40x and 63x. Images were processed with Image J software.

### Puromycin incorporation assay

Cells were seeded in 3cm dishes (4 x 10^4^) and allowed to adhere overnight before treated with 100ng/ml Dox for 2 days to knockdown expression of the endogenous eIF3d. Puromycin (A1113803, Gibco™ Puromycin Dihydrochloride) was added to cells for 10 min at a final concentration of 10 μg/ml prior to harvesting and lysis. The anti-puromycin antibody (MABE343, Merck) was used at a dilution of 1:25,000.

### Cell viability assay

The lentiviral cell lines were seeded at a density of 6,000 cells per well in 24-well plates and allowed to adhere overnight before being treated with 100 ng/mL doxycycline for 2 days. Cell viability was evaluated using the Cell Counting Kit-8 (CCK8, 7368, Tocris) according to the manufacturer’s instructions. Briefly, the cells were incubated with 10% CCK8 solution at 37 °C for 2 hours. Then, 100 µL of the culture medium was transferred to 96-well plates, and the absorbance was measured at 450 nm using a plate reader (SPECTROstar Nano).

### *C. elegans* RNAi and heat shock

*C. elegans* were fed *Escherichia coli* strain OP50 and maintained at 20 °C on nematode growth medium (NGM) composed of 50 mM NaCl, 0.25% (w/v) Bacto-peptone, and 1.7% (w/v) agar, supplemented with 1 mM MgSO₄, 1 mM CaCl₂, 25 mM KH₂PO₄ (pH 6.0), and 5 µg/mL cholesterol. The strains used in this study included Bristol N2 (wild-type), N2; uqEx41[Pmyo-2::venus::tiar-2], and N2; uqIs24[Pmyo-2::tagRFP::pab-1], kindly provided by Della David (German Center for Neurodegenerative Diseases, Tübingen). Gravid worms were synchronised in bleaching solution (0.7M NaOH, 20% (v/v) sodium hypochlorite) on the NGM plate before the stress response experiments. RNAi plasmid containing *E. coli* were obtained from the Ahringer or Vidal RNAi libraries [76, 77]. All *E. coli* HT115 (DE3) strains harboring RNAi constructs were confirmed by sequencing (Eurofins Scientific). RNAi by feeding was performed as previously described [76]. Briefly, NGM plates were supplemented with Carbenicillin (50 µg/ml) (1273-7149, ThermoFisher scientific), IPTG (1 mM) (11858905, Promega). Worms were randomly allocated to either control plates with HT115 (DE3) *E. coli* bacteria transformed with empty L4440 plasmid vector or to plates with HT115 (DE3) *E. coli* expressing the RNAi. Worms were synchronised in bleaching solution on plates and fed with RNAi bacteria until they grew to day 1 adults. The day 1 adult worms were placed in a 36℃ water bath for 3h prior to imaging.

### *C. elegans* imaging

For live imaging, a maximum of 15 worms were paralysed with 20mM tetramisole for 2-3 min on a 2% agarose pad [0.2g agarose in 20 ml M9 buffer (42mM Na2HPO4.12H2O, 22mM KH2PO4 (pH 6), 86mM NaCl, 1mM MgSO4)] before observation. A Leica DM500 B Fluorescence microscope was used with filter sets for Texas Red and FITC. Images were collected with a Leica DCF340 FX camera at a magnification of 40x and an exposure of 168ms using LeicaSuite v4.0 software. The brightfield images were taken using DIC microscopy with an exposure of 40ms.Images were processed with Image J software. The representative and quantificational images of RNAi experiments presented here were obtained using a Dragonfly upright confocal using a 40x Plan Fluotar objective. Each *C. elegans* pharynx was selected as the region of interest and stress granules were segmented from the image background using a threshold range of 50–132 GV. A particle analysis script in ImageJ was then applied to quantify stress granule sizes. To minimize the influence of imaging artifacts and background noise, granules larger than 1 µm² were outlined. The total area of these granules in each worm was subsequently used to compare differences between experimental conditions.

### Statistical analysis

For immunofluorescence experiments, statistical analysis was performed on three biological repeats unless otherwise stated. 100 cells were counted in each repeat and the analysis performed using PRISM software and one or two-way ANOVA as stated in the figure legends. For puromycin incorporation assays, quantification was performed by measuring the intensity of the puromycin signal using Image Studio Lite software in each of five biological repeats. Puromycin signal normalisation for each sample was performed against the Coomassie Blue intensity. Statistical analysis by one-way ANOVA or t-test was performed using PRISM software. For *C. elegans*, at least 30 worms were analysed per condition across three biological replicates and the area of TIAR-2–positive granules per worm quantified using Image J software. Statistical analysis by two-way ANOVA was performed using PRISM software.

## DATA AVAILABILITY

Reagents are available upon request to the corresponding author.

## ACKNOWLEDGEMENTS

We thank Della David (The Babraham Institute, Cambridge, UK) for providing the reporter strains and Laurent Désaubry for providing FL3. We thank Weitao Xiao for technical help. This work was supported by Biotechnology and Biological Sciences Research Council (BBSRC) project grants BB/V015109/1 to M.P.A and BB/X006859/1 to G.B.P. A.P.S was funded as part of a Wellcome Trust Ph.D studentship programme.

## AUTHOR CONTRIBUTION

AJW, MPA and GBP designed the study with assistance from JL and APS. JL, APS, PB and AJW performed the experiments. WZ and CT developed and assisted with methodology. AJW and JL wrote the paper. All authors read, helped revise and approved the final paper.

## ETHICS STATEMENT

Study did not require ethical approval

