## Supplementary Figures for "Eukaryotic translation initiation factor 3d regulates stress granule assembly via its RNA binding domain"

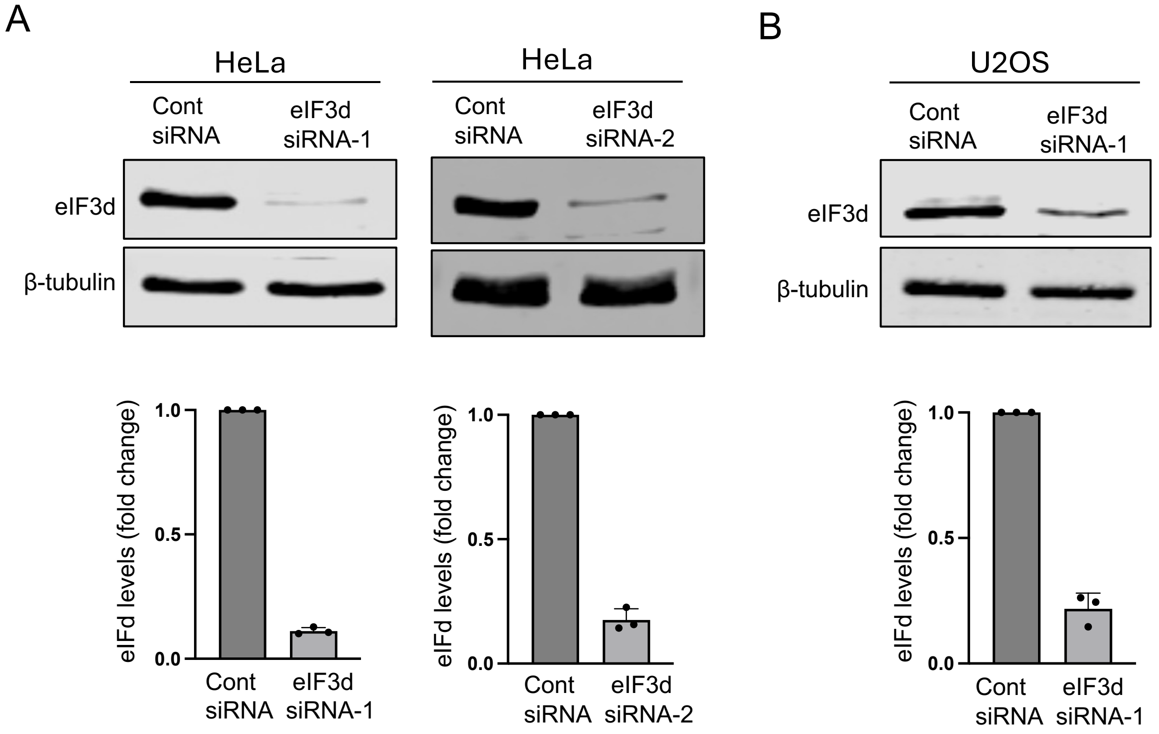


**Supplementary Figure 1. Verifying knock-down efficiency of eIF3d siRNAs. A**. HeLa cells were transfected with two siRNAs targeting eIF3d (siRNA-1 and siRNA-2) and knockdown efficiency determined by immunoblot with an eIF3d antibody. **B**. U2OS cells were transfected with control (Cont) siRNA or eIF3d siRNA-1 and eIF3d levels assessed by immunoblot. An antibody against β-tubulin was used to indicate equal loading of lysates on the gels. Quantifications of the level of eIF3d knockdown are shown beneath the blots.

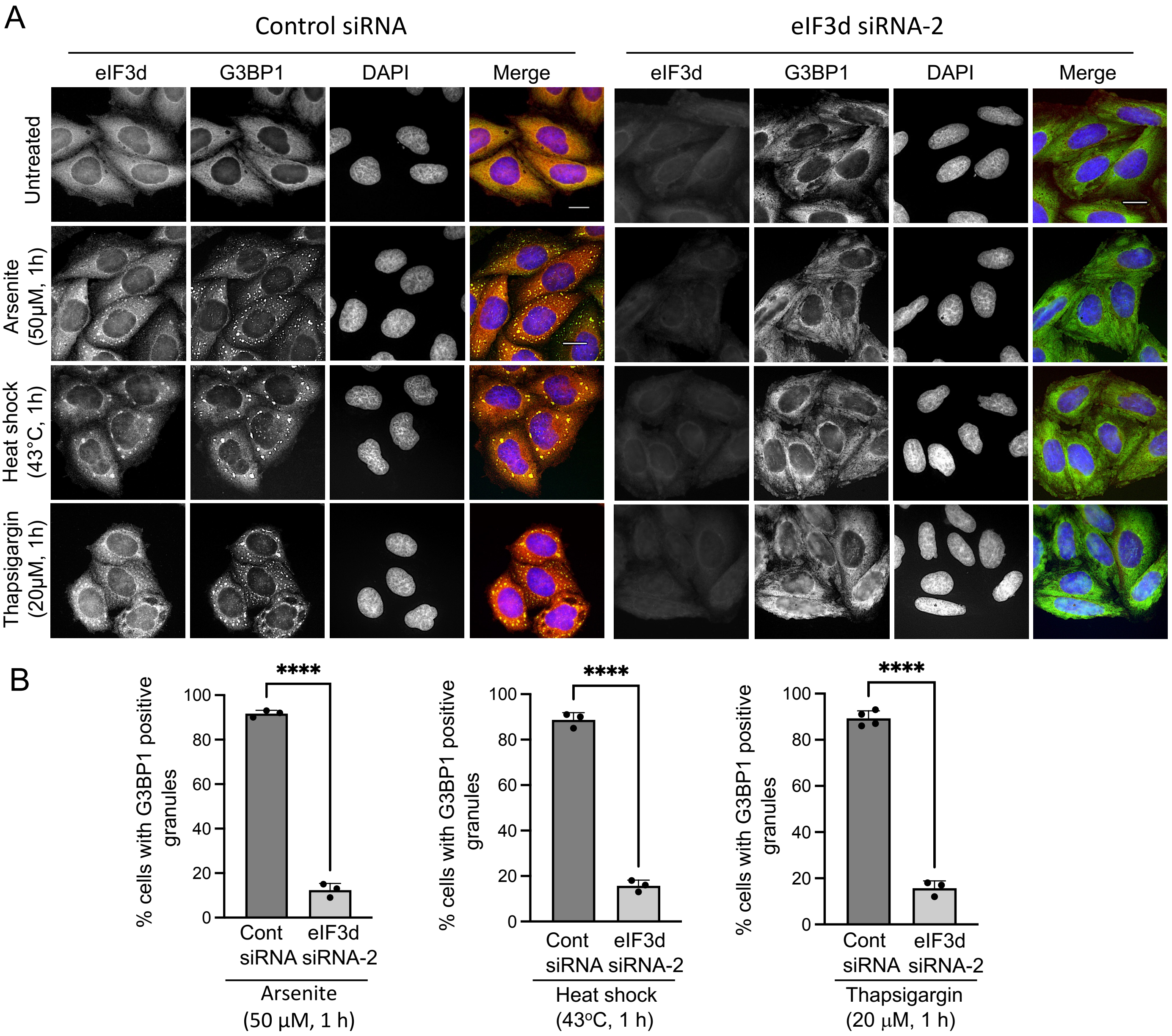


**Supplementary Figure 2. Confirming the effect of eIF3d knockdown on SG assembly in HeLa. A.** Knock-down of eIF3d with siRNA-2 blocks SG formation in HeLa cells in response to heat and ER stress. Cells were transfected with either control (Cont) siRNA or eIF3d siRNA-2 for 48 h before treatment with the indicated stresses. Cells were probed with antibodies against eIF3d and G3BP1. Nuclei were stained with DAPI. Representative images of cells are shown. Scale bar = 20 µm. **B**. Quantification of cells with G3BP1-positive granules under the indicated conditions. A total of 100 cells were analysed in each of three biological replicates. Statistical analysis was performed using unpaired t-tests (*ns* = not significant, ****P < 0.0001).

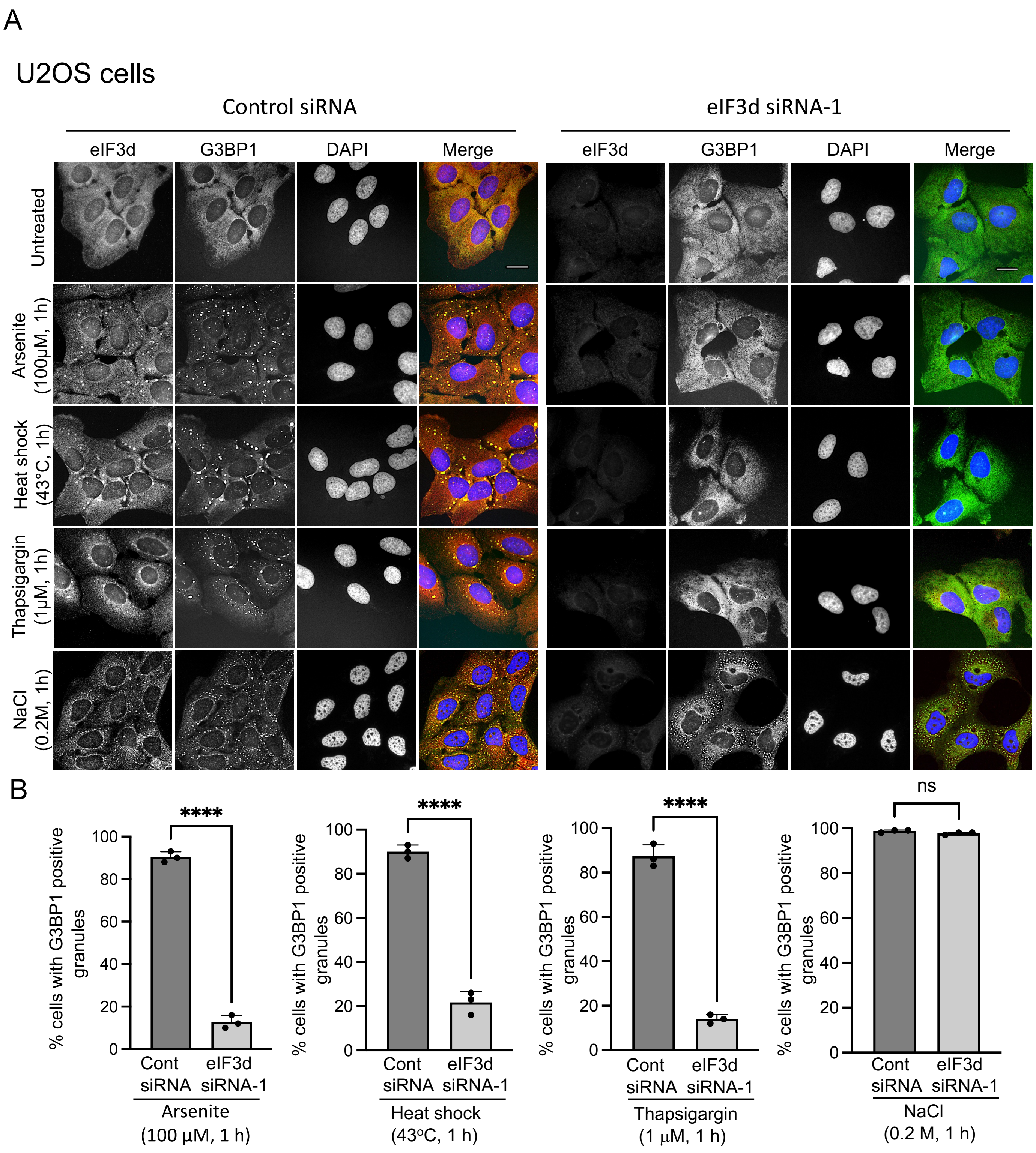

A
U2OS cells
Control siRNA
eIF3d siRNA-1
eIF3d
G3BP1
DAPI
Merge
Untreated
Arsenite (100μM, 1h)
Heat shock (43°C, 1h)
Thapsigargin (1μM, 1h)
NaCl (0.2M, 1h)
B
% cells with G3BP1 positive granules
****
ns
0
20
40
60
80
100
Cont siRNA
eIF3d siRNA-1
Arsenite (100 μM, 1 h)
Heat shock (43°C, 1 h)
Thapsigargin (1 μM, 1 h)
NaCl (0.2 M, 1 h)

**Supplementary Figure 3. eIF3d is required for SG assembly in U2OS. A.** Knock-down of eIF3d blocks SG formation in U2OS cells in response to the indicated stresses. Cells were transfected with either control (Cont) siRNA or eIF3d siRNA-1 for 48 h before treatment with the indicated stresses. Cells were probed with antibodies against eIF3d and G3BP1. Nuclei were stained with DAPI. Representative images of cells are shown. Scale bar = 20 µm. **B**. Quantification of cells with G3BP1-positive granules under the indicated conditions. A total of 100 cells were analysed in each of three biological replicates. Statistical analysis was performed using unpaired t-tests (*ns* = not significant, ****P < 0.0001).

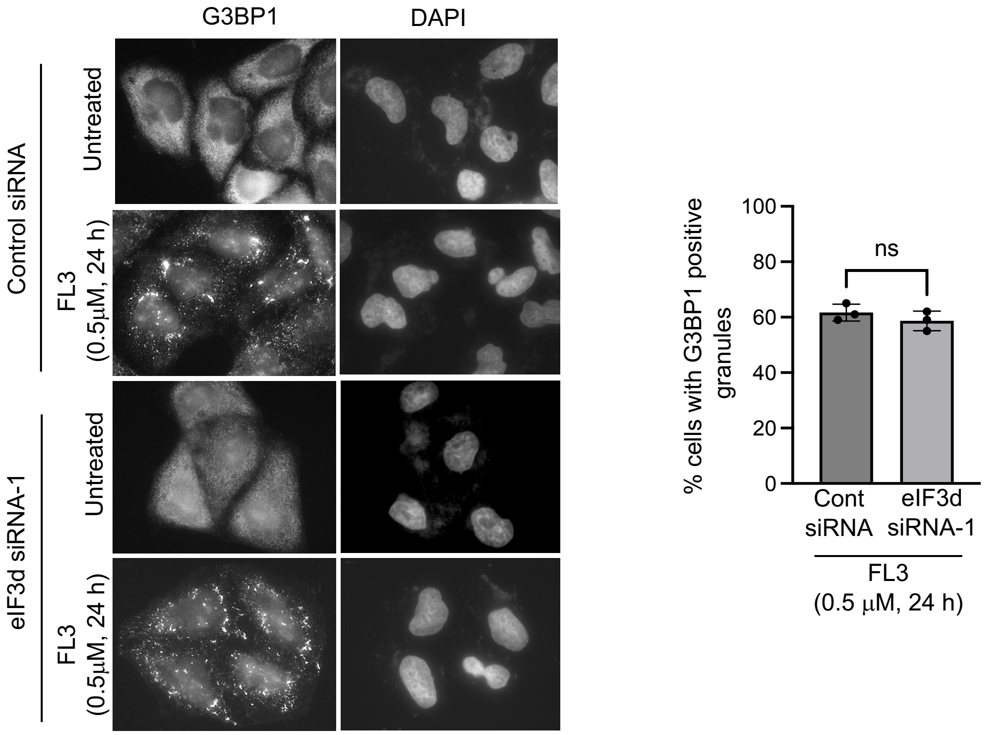


**Supplementary Figure 4. eIF3d is not required for SG assembly in response to eIF4A inhibition by FL3.** Cells were transfected with either control (Cont) siRNA or eIF3d siRNA-1 for 48 h before treatment with 0.5 μM FL3 for 24 h. Cells were probed with antibodies against G3BP1. Nuclei were stained with DAPI. Representative images of cells are shown along with quantification of cells with G3BP1-positive granules under the indicated conditions. A total of 100 cells were analysed in each of three biological replicates. Statistical analysis was performed using an unpaired t-test (*ns* = not significant).

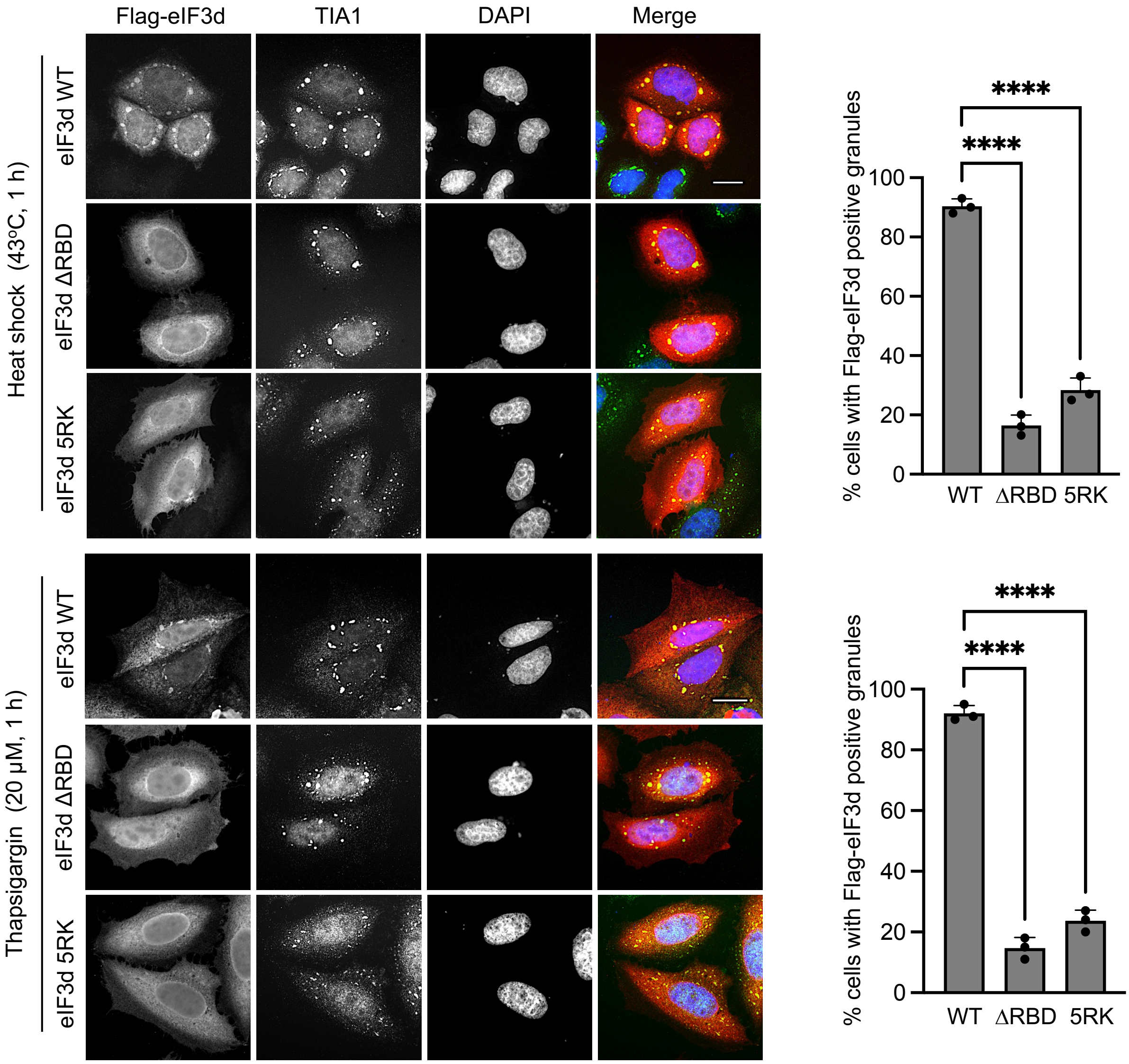


**Supplementary Figure 5. The RBD of eIF3d is required for its recruitment to SGs in response to heat and ER stress in HeLa.** Cells were transfected with plasmids expressing Flag-tagged wild-type eIF3d or the mutants eIF3d(ΔRBD) and eIF3d(5RK) before treatment with the indicated stresses. Cells were probed with antibodies against the Flag-tag and TIA1. Nuclei were stained with DAPI. Representative images of cells are shown. Scale bar = 20 µm. Quantification of cells with Flag-eIF3d-positive granules is shown on the right. A total of 100 cells were analysed in each of three biological replicates. Statistical analysis was performed using one-way ANOVA (*ns* = not significant, ****P < 0.0001).

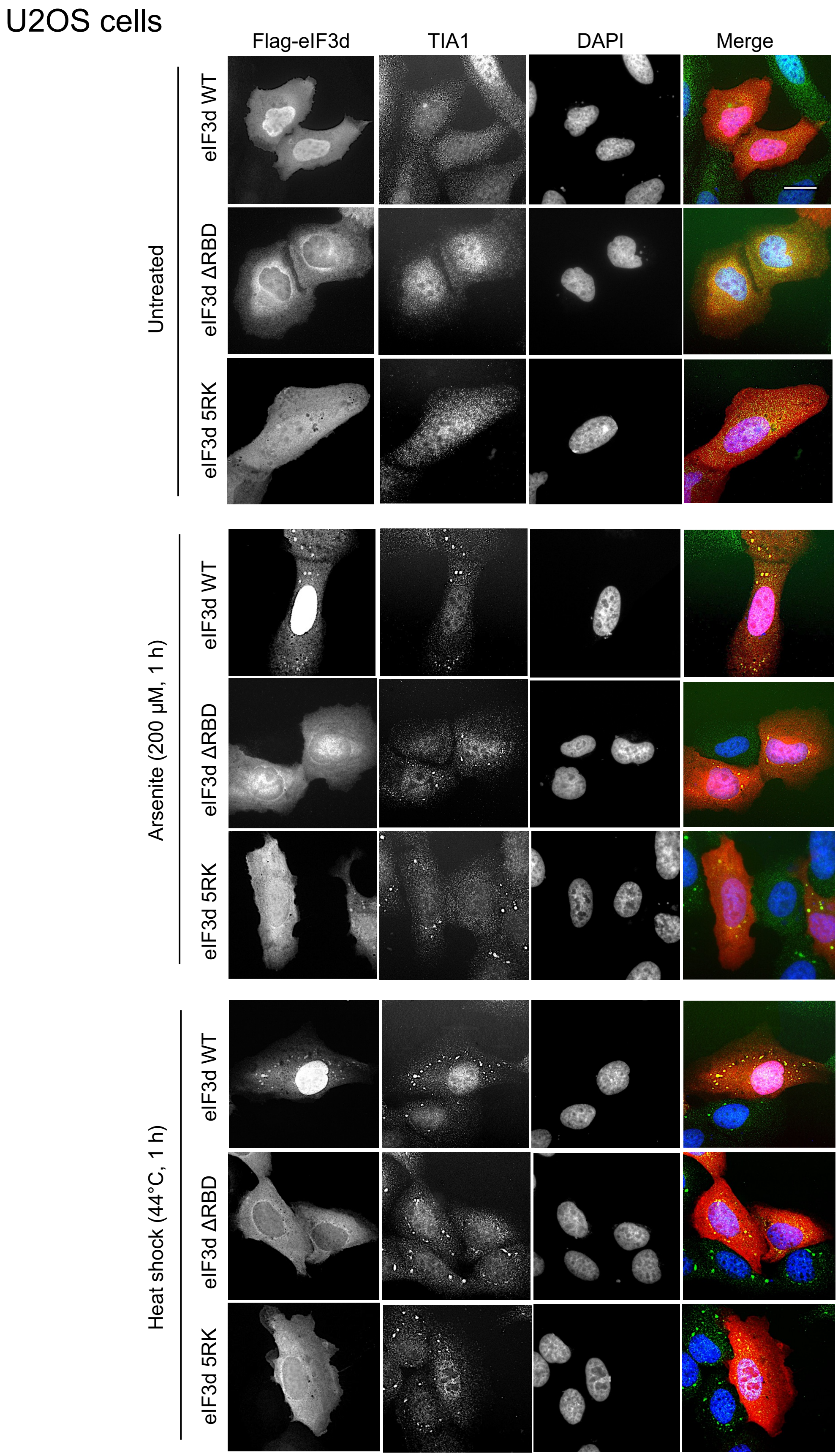
U2OS cells
Flag-eIF3d
TIA1
DAPI
Merge
Untreated
eIF3d WT
eIF3d ΔRBD
eIF3d 5RK
Arsenite (200 µM, 1 h)
eIF3d WT
eIF3d ΔRBD
eIF3d 5RK
Heat shock (44°C, 1 h)
eIF3d WT
eIF3d ΔRBD
eIF3d 5RK

**Supplementary Figure 6. The RBD of eIF3d is required for its recruitment to SGs in response to stress in U2OS.** Cells were transfected with plasmids expressing Flag-tagged wild-type eIF3d or the mutants eIF3d(ΔRBD) and eIF3d(5RK) before treatment with the indicated stresses. Cells were probed with antibodies against the Flag-tag and TIA1. Nuclei were stained with DAPI. Representative images of cells are shown. Scale bar = 20 μm.

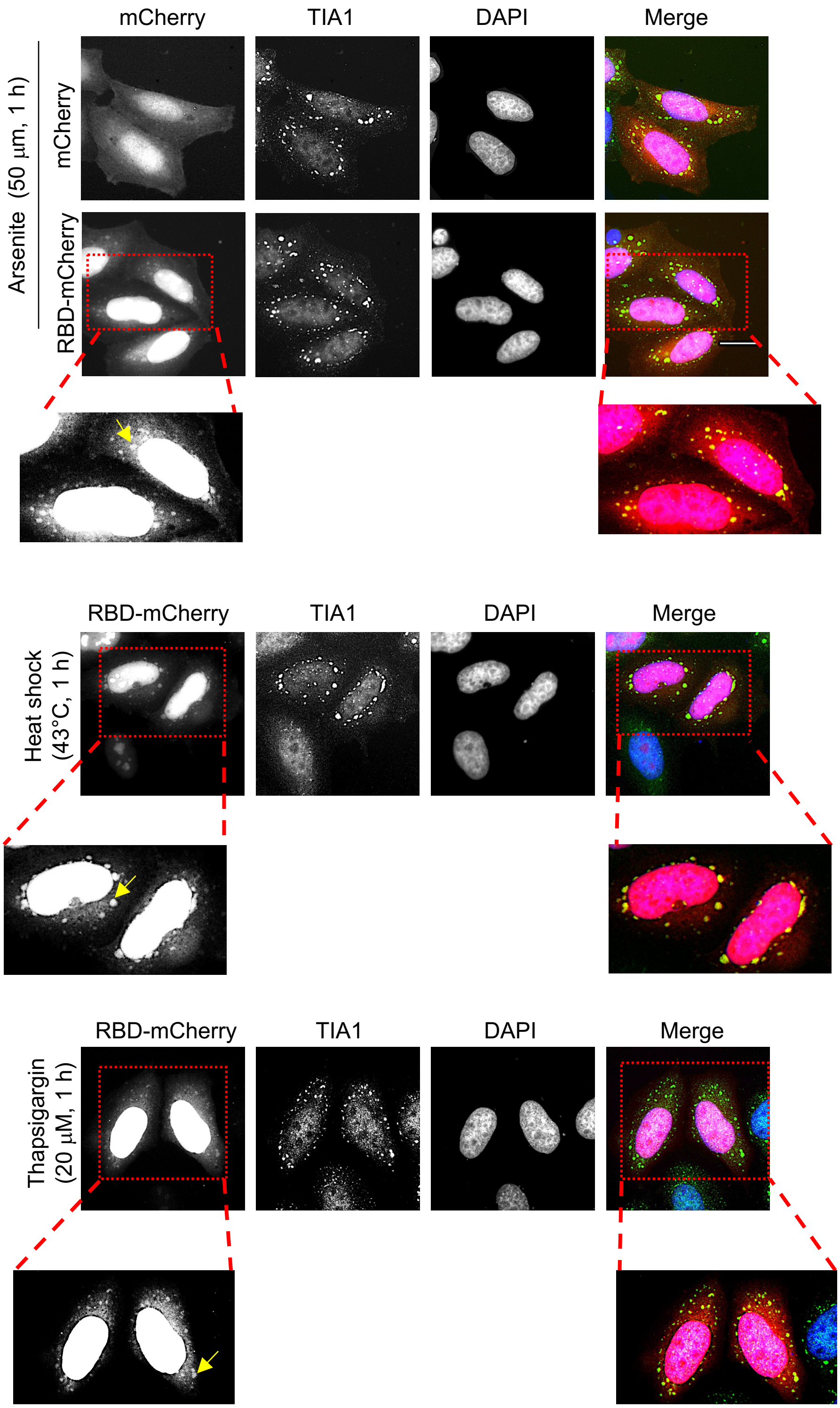


**Supplementary Figure 7. The RBD of eIF3d is recruited to SGs in response to stress.** HeLa cells were transfected with plasmids expressing either mCherry or the eIF3d RBD fused to mCherry and treated as indicated. TIA1 immunofluorescence was used to visualize SGs. Scale bar = 20 µm. Expanded overexposed images are shown to visualise RBD-mCherry in SGs. Arrows indicate SGs.

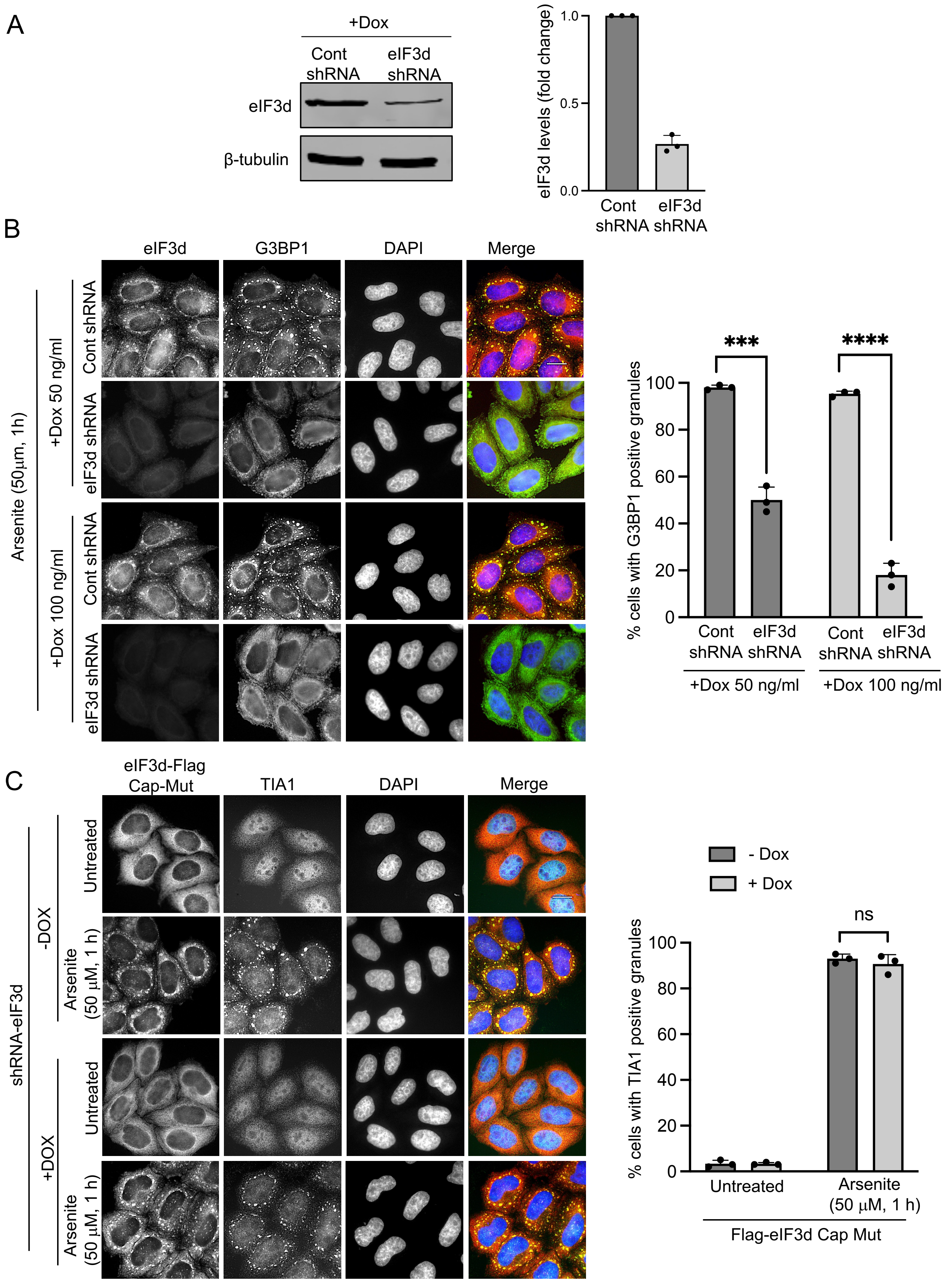

A
+Dox
Cont shRNA
eIF3d shRNA
eIF3d
β-tubulin
eIF3d levels (fold change)
1.0
0.5
0.0
Cont shRNA
eIF3d shRNA
B
eIF3d
G3BP1
DAPI
Merge
Arsenite (50μm, 1h)
+Dox 50 ng/ml
+Dox 100 ng/ml
Cont shRNA
eIF3d shRNA
% cells with G3BP1 positive granules
***
****
100
80
60
40
20
0
Cont shRNA
eIF3d shRNA
+Dox 50 ng/ml
+Dox 100 ng/ml
C
eIF3d-Flag Cap-Mut
TIA1
DAPI
Merge
shRNA-eIF3d
-DOX
+DOX
Untreated
Arsenite (50 μM, 1 h)
- Dox
+ Dox
ns
% cells with TIA1 positive granules
Untreated
Arsenite (50 μM, 1 h)
Flag-eIF3d Cap Mut

**Supplementary Figure 8. Doxycycline-inducible knockdown of eIF3d blocks SG assembly in HeLa and this can be rescued by an eIF3d cap-binding mutant. A.** Stable HeLa cells featuring a doxycycline (Dox)-inducible control (Cont) shRNA or eIF3D shRNA were treated with Dox and the levels of eIF3d determined by immunoblot. An antibody targeting β-tubulin was used to indicate equal loading of lysates on the gels. **B**. The control-shRNA and eIF3D-shRNA cells were subjected to the indicated concentrations of Dox and treated with 50 μM sodium arsenite for 1 hr. The cells were probed with eIF3d and G3BP1 antibodies. Nuclei were stained with DAPI. Representative images of cells are shown. Scale bar = 20 μm. Quantification of cells with G3BP1-positive granules is shown on the right. A total of 100 cells were analysed in each of three biological replicates. Statistical analysis was performed using an unpaired t-test (*ns* = not significant, ***P = 0.0001 ****P = <0.0001). **C.** eIF3d-shRNA cells stably expressing a Flag-tagged cap-binding mutant of eIF3d were treated with or without 100 ng/ml Dox to induce shRNA expression. The cells were then treated with 50 μM sodium arsenite for 1 hr and probed with anti-Flag and TIA1 antibodies. Nuclei are stained with DAPI. Representative images of cells are shown. Scale bar = 20 μm. Quantification of cells with TIA1-positive granules is shown on the right. A total of 100 cells were analysed in each of three biological replicates. Statistical analysis was performed using an unpaired t-test (*ns* = not significant).

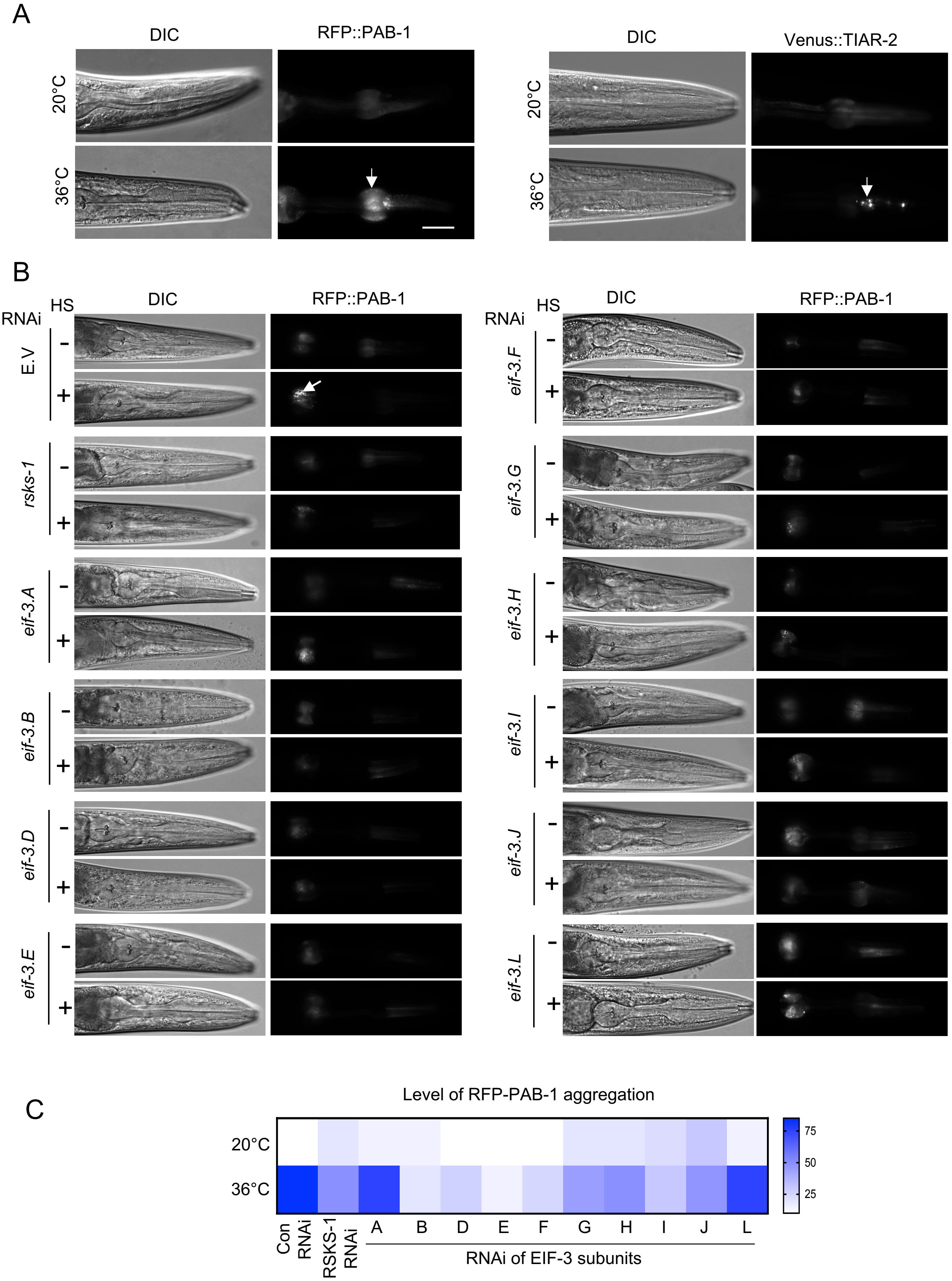
A
DIC
RFP::PAB-1
DIC
Venus::TIAR-2
20°C
36°C
B
RNAi
HS
DIC
RFP::PAB-1
E.V
rsks-1
eif-3.A
eif-3.B
eif-3.D
eif-3.E
eif-3.F
eif-3.G
eif-3.H
eif-3.I
eif-3.J
eif-3.L
-
+
C
Level of RFP-PAB-1 aggregation
20°C
36°C
Con RNAi
RSKS-1 RNAi
A
B
D
E
F
G
H
I
J
L
RNAi of EIF-3 subunits
75
50
25

**Supplementary Figure 9. RNAi screen in *C. elegans* to determine importance of EIF-3 factors in SG assembly**. **A**. Worms expressing RFP::PAB-1 and Venus::TIAR-2 fluorescent reporter genes in the pharynx. DIC and fluorescent images are presented. Application of heat stress (HS; 36$^{o}$C, 3 hrs) induces RFP::PAB-1 and Venus-TIAR-2 assembly into granules. **B**. RNAi screen of EIF-3 factors using the RFP::PAB-1 reporter strain as a readout of granule assembly. DIC and fluorescent images are presented. An RNAi targeting the worm orthologue of human S6 kinases, RSKS-1, was used as a positive control [33]. Arrows indicate granules. Scale bar = 25 µm. **C.** Heat map showing RNAi screening results for different EIF-3 subunits based on RFP::PAB-1 aggregation levels in the pharynx. The proportion of worms with PAB-1-positive granules following RNAi feeding is represented by colour intensities. The scale represents the % of worms displaying granules for each condition. At least 45 worms were analysed per condition across three biological replicates.

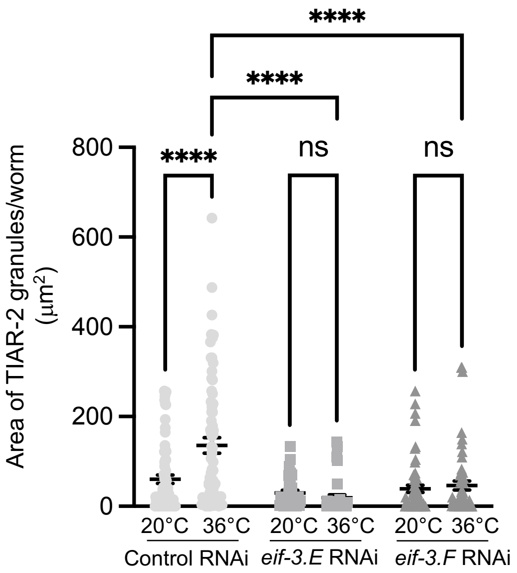


**Supplementary Figure 10. RNAi screen in *C. elegans* to determine importance of EIF-3 factors in SG assembly**. Worms expressing pharyngeal RFP::PAB-1 and Venus::TIAR-2 reporters were fed *eif-3.E* or *eif-3.F* RNAi and subjected to heat shock at 36$^{o}$C for 3 h. The total area of TIAR-2–positive granules per worm was quantified. At least 30 worms were analysed per condition across three biological replicates. Data were analysed by two-way ANOVA (*ns* = not significant; ****$p < 0.0001$).

### Figure 3C

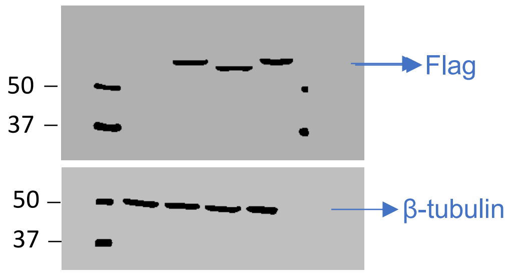


### Figure 6.B

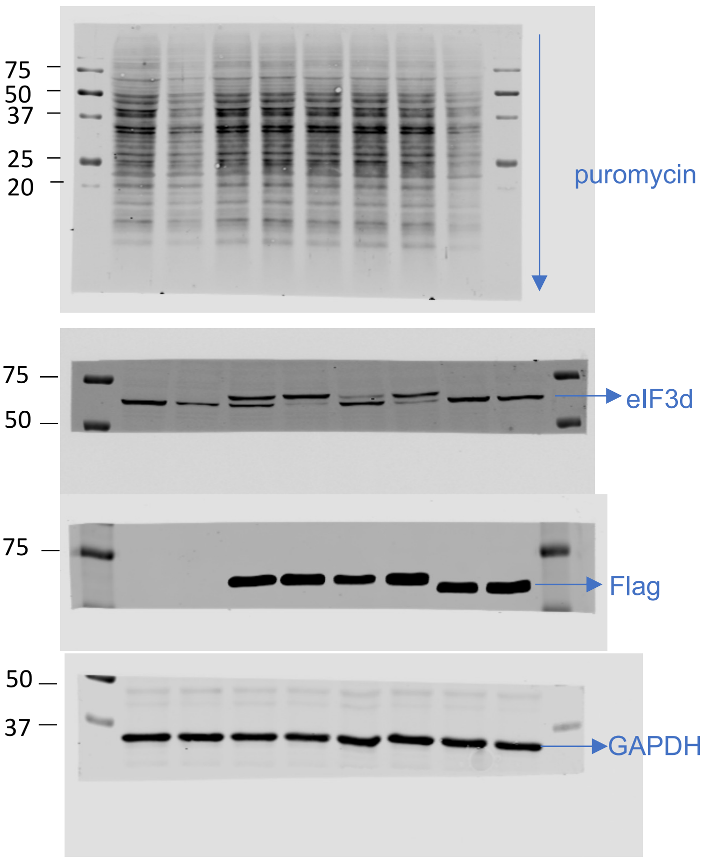


### Figure S3

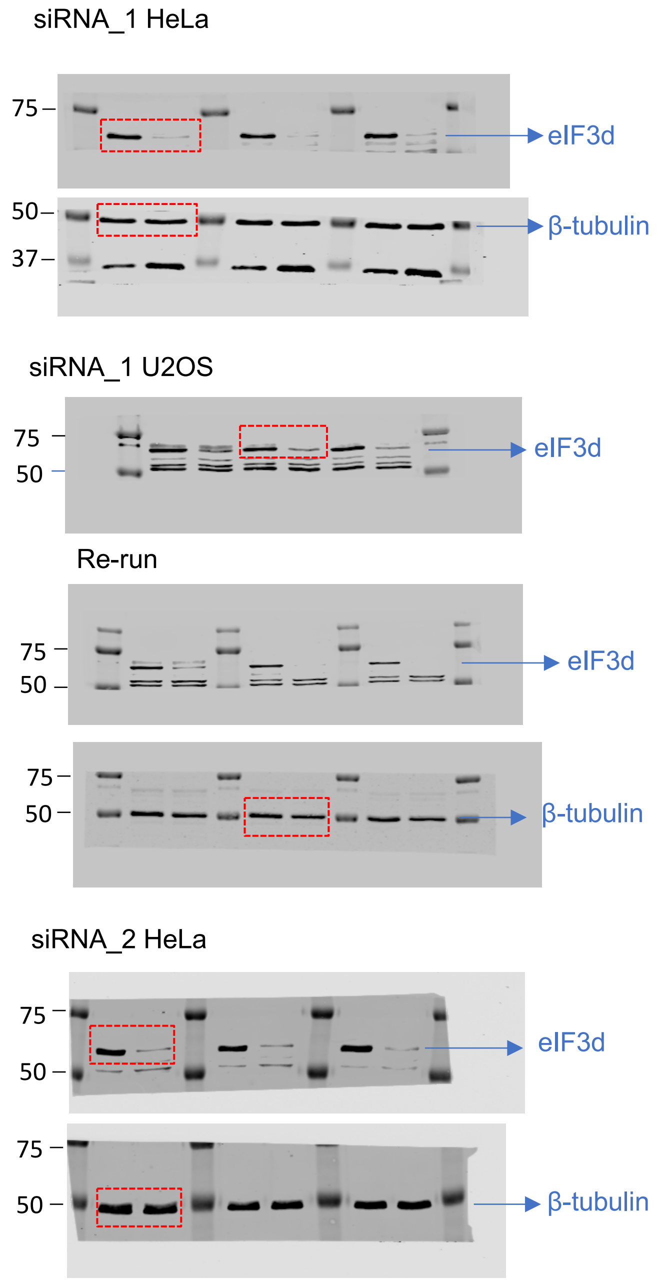


### Figure S10

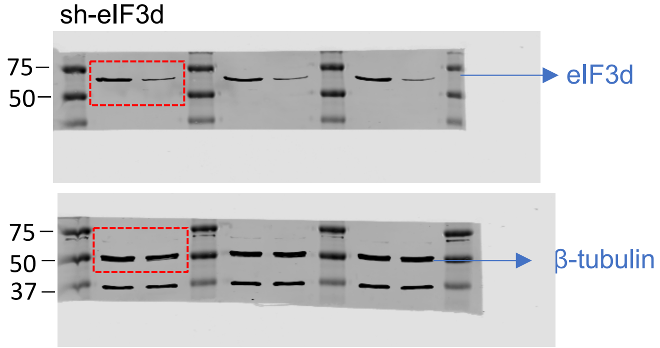


**Supplementary Figure 11. Unprocessed scans of immunoblots.**

**Table S1. Antibodies used in the study**

| Antibody | Cat Number | Company |
|---|---|---|
| Primary | | |
| eIF3d | PA5-9648 | Invitrogen |
| G3BP1 (H-10) | SC-36533 | Santa Cruz |
| TIA1 | AB140595 | Abcam |
| Anti-Flag M2 | F1804 | Sigma-Aldrich |
| β-Tubulin | ab6046 | Abcam |
| GAPDH | AC001 | Abclonal |
| Puromycin | MABE343 | Merck |
| Secondary WB antibody | | |
| IRDye® 800CW anti-Rabbit | 926-32213 | LI-COR |
| IRDye® 800CW anti-Mouse | 926-32212 | LI-COR |
| Secondary IF antibody | | |
| Alexa Fluor™ Plus 594 anti-Rabbit | A32754 | Invitrogen |
| Alexa Fluor™ Plus 488 anti-Mouse | A32723 | Invitrogen |
| Alexa Fluor™ Plus 594 anti-Mouse | A32742 | Invitrogen |
| Alexa Fluor™ Plus 488 anti-Rabbit | A11008 | Invitrogen |
| Prolong® Diamond-DAPI | P36962 | ThermoFisher |
